# Bacterial colonization stimulates a complex physiological response in the immature human intestinal epithelium

**DOI:** 10.1101/144568

**Authors:** David R. Hill, Sha Huang, Melinda S. Nagy, Veda K. Yadagiri, Courtney Fields, Dishari Mukherjee, Brooke Bons, Priya H. Dedhia, Alana M. Chin, Yu-Hwai Tsai, Shrikar Thodla, Thomas M. Schmidt, Seth Walk, Vincent B. Young, Jason R. Spence

## Abstract

The human gastrointestinal tract is immature at birth, yet must adapt to dramatic changes such as oral nutrition and microbial colonization. The confluence of these factors can lead to severe inflammatory disease in premature infants; however, investigating complex environment-host interactions is difficult due to limited access to immature human tissue. Here, we demonstrate that the epithelium of human pluripotent stem cell-derived human intestinal organoids is globally similar to the immature human epithelium and we utilize HIOs to investigate complex host-microbe interactions in this naïve epithelium. Our findings demonstrate that the immature epithelium is intrinsically capable of establishing a stable host-microbe symbiosis. Microbial colonization leads to complex contact and hypoxia driven responses resulting in increased antimicrobial peptide production, maturation of the mucus layer, and improved barrier function. These studies lay the groundwork for an improved mechanistic understanding of how colonization influences development of the immature human intestine.

## Introduction

The epithelium of the gastrointestinal (GI) tract represents a large surface area for host-microbe interaction and mediates the balance between tolerance of mutualistic organisms and the exclusion of potential pathogens (Peterson and Artis, 2014). This is accomplished, in part, through the formation of a tight physical epithelial barrier, in addition to epithelial secretion of anti-microbial peptides and mucus (Veereman-Wauters, 1996; Renz et al., 2012). Development and maturation of the epithelial barrier coincides with the first exposure of the GI tract to microorganisms and the establishment of a microbial community within the gut (Palmer et al., 2007; Koenig et al., 2011).

Although microorganisms have long been appreciated as the primary drivers of the postnatal expansion of adaptive immunity (Renz et al., 2012; Shaw et al., 2010; Hviid et al., 2011; Abrahamsson et al., 2014; Arrieta et al., 2015), and more recently as key stimuli in the development of digestion (Erkosar et al., 2015), metabolism (Cho et al., 2012), and neurocognitive function (Diaz Heijtz et al., 2011; Clarke et al., 2014; Borre et al., 2014; Desbonnet et al., 2014), it remains unclear how the human epithelial surface adapts to colonization and expansion of microorganisms within the immature GI tract.

Studies in gnotobiotic mice have improved our understanding of the importance of microbes in normal gut function since these mice exhibit profound developmental defects in the intestine (Round and Mazmanian, 2009; Gensollen et al., 2016; Bry et al., 1996; Hooper et al., 1999) including decreased epithelial turnover, impaired formation of microvilli (Abrams et al., 1963), and altered mucus glycosylation at the epithelial surface (Bry et al., 1996; Goto et al., 2014; Cash et al., 2006). However, evidence also suggests that the immature human intestine may differ significantly from the murine intestine, especially in the context of disease (Nguyen et al., 2015). For example, premature infants can develop necrotizing enterocolitis (NEC), an inflammatory disease with unknown causes. Recent reports suggest a multifactorial etiology by which immature intestinal barrier function predisposes the preterm infant to intestinal injury and inflammation following postpartum microbial colonization (Neu and Walker, 2011; Morrow et al., 2013; Greenwood et al., 2014; Hackam et al., 2013; Afrazi et al., 2014; Fusunyan et al., 2001; Nanthakumar et al., 2011). Rodent models of NEC have proven to be inadequate surrogates for studying human disease (Tanner et al., 2015). Therefore, direct studies of host-microbial interactions in the immature human intestine will be important to understand the complex interactions during bacterial colonization that lead to a normal gut development or disease.

Important ethical and practical considerations have limited research on the immature human intestine. For example, neonatal surgical specimens are often severely damaged by disease and not conducive for *ex vivo* studies. We and others have previously demonstrated that human pluripotent stem cell derived human intestinal organoids (HIOs) closely resemble immature intestinal tissue (Spence et al., 2011; ***Finkbeiner et al., 2015b***; Watson et al., 2014; Forster et al., 2014; Dedhia et al., 2016; Aurora and Spence, 2016; Chin et al., 2017) and recent work has established gastrointestinal organoids as a powerful model of microbial pathogenesis at the mucosal interface (Leslie et al., 2015; McCracken et al., 2014; Forbester et al., 2015; Hill and Spence, 2017).

In the current work, we used HIOs as a model immature intestinal epithelium and a human-derived non-pathogenic strain of *E. coli* as a model intestinal colonizer to examine how host-microbe interactions affected intestinal maturation and function. Although the composition of the neonatal intestinal microbiome varies between individuals, organisms within the genera *Escherichia* are dominant early colonizers (Gosalbes et al., 2013; Bäckhed et al., 2015) and non-pathogenic *E. coli* are widely prevalent and highly abundant components of the neonatal stool microbiome (Palmer et al., 2007; Koenig et al., 2011; Bäckhed et al., 2015; Morrow et al., 2013). Microinjection of *E. coli* into the lumen of 3-dimensional HIOs resulted in stable bacterial colonization *in vitro*, and using RNA-sequencing, we monitored the global transcriptional changes in response to colonization. We observed widespread, time-dependent transcriptional responses that are the result of both bacterial contact and luminal hypoxia resulting from bacterial colonization in the HIO. Bacterial association with the immature epithelium increased antimicrobial defenses and resulted in enhanced epithelial barrier function and integrity. We observed that NF-*κ*B is a central downstream mediator of the transcriptional changes induced by both bacterial contact and hypoxia. We further probed the bacterial contact and hypoxia dependent epithelial responses using experimental hypoxia and pharmacological NF-*κ*B inhibition, which allowed us to delineate which of the transcriptional and functional responses of the immature epithelium were oxygen and/or NF-*κ*B dependent. We found that NF-*κ*B dependent microbe-epithelial interactions were beneficial by enhancing barrier function and protecting the epithelium from damage by inflammatory cytokines. Collectively, these studies shed light on how microbial contact with the immature human intestinal epithelium can lead to modified function.

## Results

### Pluripotent stem-cell derived intestinal epithelium transcriptionally resembles the immature human intestinal epithelium

Previous work has demonstrated that stem cell derived human intestinal organoids resemble immature human duodenum (Watson et al., 2014; ***Finkbeiner et al., 2015b***; Tsai et al., 2017). Moreover, transplantation into immunocompromised mice results in HIO maturation to an adult-like state (Watson et al., 2014; ***Finkbeiner et al., 2015b***). These analyses compared HIOs consisting of epithelium and mesenchyme to whole-thickness human intestinal tissue, which also possessed cellular constituents lacking in HIOs such as neurons, blood vessels and immune cells (***Finkbeiner et al., 2015b***). Thus the extent to which the HIO epithelium resembles immature/fetal intestinal epithelium remains unclear. To address this gap and further characterize the HIO epithelium relative to fetal and adult duodenal epithelium, we isolated and cultured epithelium from HIOs grown entirely *in vitro*, from fetal duodenum, adult duodenum, or HIOs that had been transplanted into the kidney capsule of NSG immuno-deficient mice and matured for 10 weeks. These epithelium-only derived organoids were expanded *in vitro* in uniform tissue culture conditions for 4-5 passages and processed for RNA-sequencing (RNA-seq) (**Figure 1 - Supplement 1**). Comparison of global transcriptomes between all samples in addition to human embryonic stem cells (hESCs) used to generate HIOs (***Finkbeiner et al. 2015b***; E-MTAB-3158) revealed a clear hierarchy in which both *in vitro* grown HIO epithelium (*P* = 5.06 × 10^−9^) and transplanted epithelium (*P* = 7.79 × 10^−14^) shares a substantially greater degree of similarity to fetal small intestinal epithelium (**Figure 1 - Supplement 1A**).

While unbiased clustering demonstrated that transplanted epithelium is closely resembles fetal epithelium, we noted a shift towards the adult transcriptome that resulted in a relative increase in the correlation between transplanted HIO epithelium and adult duodenum-derived epithelium grown *in vitro* (**Figure 1 - Supplement 1B**, *P* = 1.17 × 10^−4^). Principle component analysis (PCA) of this multi-dimensional gene expression dataset (**Figure 1 - Supplement 1C**) corroborated the correlation analysis, and indicated that PC1 was correlated with developmental stage (PC1, 27.75% cumulative variance) and PC2 was correlated with tissue maturation status (PC2, 21.49% cumulative variance); cumulatively, PC1 and PC2 accounted for 49.24% of the cumulative variance between samples, suggesting that developmental stage and tissue maturation status are major sources of the transcriptional variation between samples. HIO epithelium clustered with fetal epithelium along PC2 whereas transplanted HIO epithelium clustered with adult epithelium.

We further used differential expression analysis to demonstrate that in vitro grown HIO epithelium is similar to the immature human intestine whereas *in vivo* transplanted HIO epithelium is similar to the adult epithelium. To do this, we identified differentially expressed genes through two independent comparisons: 1) human fetal vs. adult epithelium; 2) HIO epithelium vs. transplanted HIO epithelium. Genes enriched in transplanted HIO epithelium relative to the HIO epithelium were compared to genes enriched in the adult duodenum relative to fetal duodenum (**Figure 1 - Supplement 1D**). There was a highly significant correlation between log_2_-transformed expression ratios where transplanted HIOs and adult epithelium shared enriched genes while HIO and fetal epithelium shared enriched genes (*P* = 2.6 × 10^−28^). This analysis supports previously published data indicating that the epithelium from HIOs grown *in vitro* recapitulates the gene expression signature of the immature duodenum and demonstrates that the HIO epithelium is capable of adopting a transcriptional signature that more strongly resembles adult duodenum following transplantation into mice.

### HIOs can be stably associated with non-pathogenic *E. coli*

Given that the HIO epithelium recapitulates many of the features of the immature intestinal epithelium, we set out to evaluate the effect of bacterial colonization on the naïve HIO epithelium. Previous studies have established that pluripotent stem cell derived intestinal organoids can be injected with live viral (Finkbeiner et al., 2012) or bacterial pathogens (Leslie et al., 2015; Engevik et al., 2015; Forbester et al., 2015), however it was not known if HIOs could be stably co-cultured with non-pathogenic microorganisms. We co-cultured HIOs with the non-motile human-derived *Esherichia coli* strain ECOR2 (Ochman and Selander, 1984). Whole genome sequencing and phylogentic analysis demonstrated that *E. coli* str. ECOR2 is closely related to other non-pathogenic human *E. coli* and only distantly related to pathogenic *E. coli* and *Shigella* isolates (**Figure 1 - Supplement 3**). We developed a microinjection technique to introduce live *E. coli* into the HIO lumen in a manner that prevented contamination of the surrounding media (**Figure 1 - Supplement 2**). HIOs microinjected with 10^5^ live *E. coli* constitutively expressing GFP exhibit robust green fluorescence within 3 h of microinjection (**Figure 1A** and **Video 1**). Numerous *E. coli* localized to the luminal space at 48 h post-microinjection and are present adjacent to the HIO epithelium, with some apparently residing in close opposition to the apical epithelial surface (**Figure 1B**).

**Figure 1:**
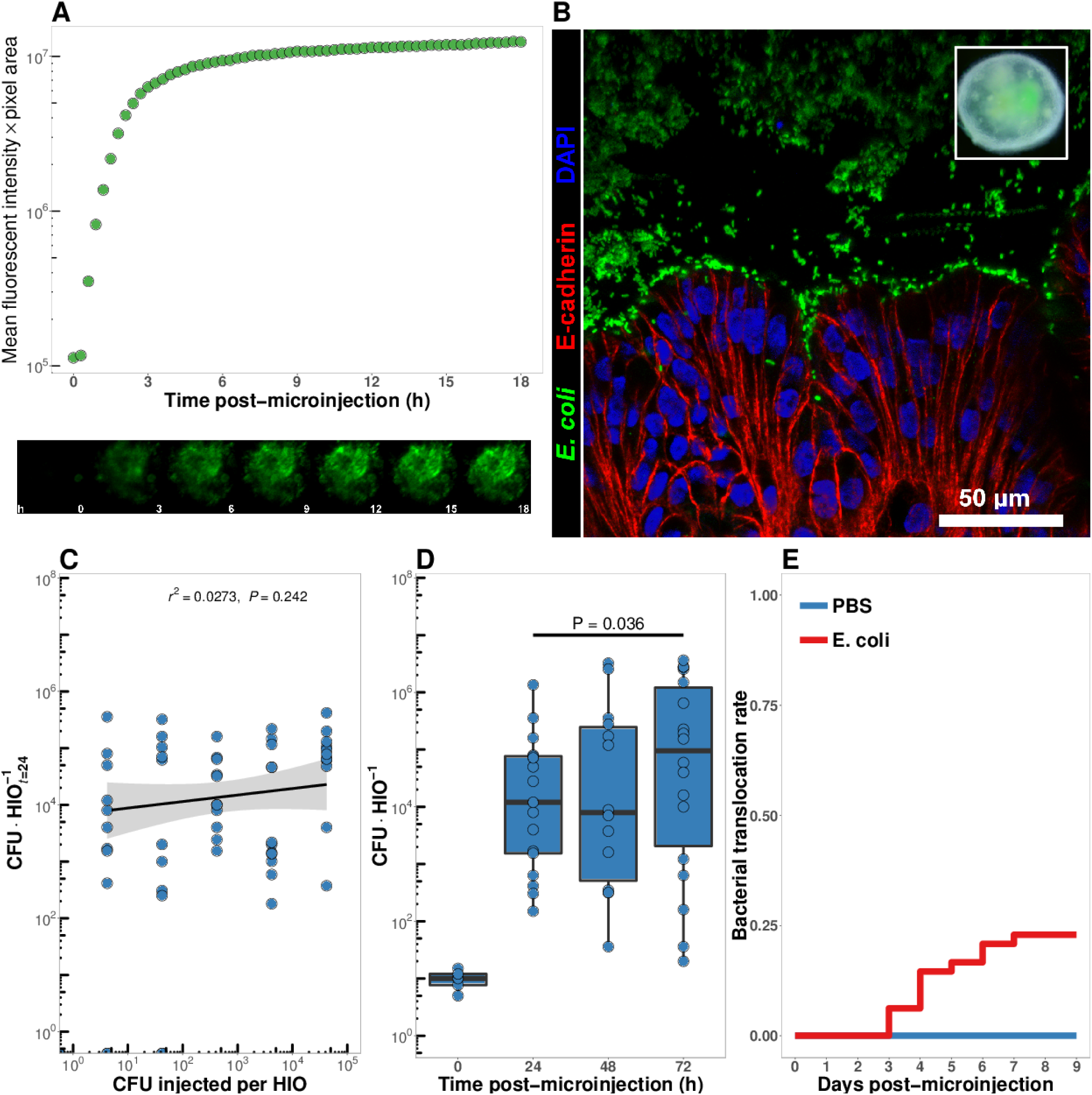
**A** Mean fluorescent intensity of a human intestinal organoid (HIO) containing live GFP^+^ *E. coli* str. ECOR2. The lower panels show representative images from the time series. Representative of 3 independent experiments. Video 1 is an animation corresponding to this dataset. **B** Confocal micrograph of the HIO epithelium (E-cadherin) in direct association with GFP+ *E. coli* at 48 h post-microinjection with 10^4^ live *E. coli*. 60X magnification. **C** Luminal CFU per HIO *E. coli* at 24 h post-microinjection relative to the injected concentration of 5 × 10^−1^ to 5 × 10^5^ CFU per HIO at the start of the experiment. *N* = 10 biological replicates per *E. coli* dose. The *r*^2^ and *P* value shown in the figure represent the results of a linear regression analysis of the relationship between the total CFU/HIO at 24 h and the initial number of CFU injected **D** Luminal CFU per HIO at 0-72 h following microinjection with 10 CFU *E. coli* per HIO. *N* = 13 - 17 replicate HIOs per time point. The *P*-value represents the results of a two-tailed Student’s *t*-test comparing the two conditions indicated. **E** Daily proportion of HIO cultures with no culturable *E. coli* in the external media following *E. coli* microinjection (*N* = 48) or PBS microinjection (*N* = 8).

In order to determine the minimum number of colony forming units (CFU) of *E. coli* required to establish short term colonization (24 hours), we microinjected increasing numbers of live *E. coli* suspended in PBS into single HIOs and collected and determined the number of bacteria in the luminal contents at 24 h post-microinjection (**Figure 1C**). Single HIOs can be stably colonized by as few as 5 CFU *E. coli* per HIO with 77.8 % success (positive luminal culture and negative external media culture at 24 h post-injection) and 100 % success at ≥ 100 CFU per HIO (**Figure 1C**). Increasing the number of CFU *E. coli* microinjected into each HIO at *t* = 0 did result in a significant increase in the mean luminal CFU per HIO at 24 hours post-microinjection at any dose (ANOVA *P* = 0.37; **Figure 1C**). Thus, the 24 h growth rate of *E. coli* within the HIO lumen 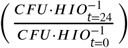 was negatively correlated with the CFU injected (r^2^ = 0.625, P = 3.1 × 10^−12^).

Next, we examined the stability of HIO and *E. coli* co-cultures over time *in vitro*. HIOs were microinjected with 10 CFU *E. coli* and maintained for 24-72 h (**Figure 1D**). Rapid expansion of *E. coli* density within the HIO lumen was observed in the first 24 h, with relatively stable bacterial density at 48-72 hr. A 6.25-fold increase in bacterial density was observed between 24 and 72 h post-microinjection (*P* = 0.036). Importantly, samples taken from the external HIO culture media were negative for *E. coli* growth.

Finally, we examined the stability of HIO cultures following *E. coli* microinjection (**Figure 1E**). A total of 48 individual HIOs were microinjected with 10^4^ CFU *E. coli* each. Controls were microinjected with sterile PBS alone. We found that external culture media was sterile in 100% of control HIOs throughout the entire experiment, and in 100 % of *E. coli* injected HIOs on days 0-2 post-microinjection. On days 3-9 post-microinjection some cultured media was positive for *E. coli* growth; however, 77.08 % of *E. coli* injected HIOs were negative for *E. coli* in the external culture media throughout the timecourse. Additional control experiments were conducted to determine if the HIO growth media had any effect on *E. coli* growth. *E.coli*-inoculated HIO growth media showed that the media itself allowed for robust bacterial growth, and therefore the absence of *E. coli* growth in external media from HIO cultures could not be attributed to the media composition alone (**Figure 1 - Supplement 3**). Thus, the large majority of *E. coli* colonized HIOs remain stable for an extended period when cultured *in vitro* and without antibiotics.

### Bacterial colonization elicits a broad-scale, time-dependent transcriptional response

Colonization of the immature gut by microbes is associated with functional maturation in both model systems(Kremer et al., 2013; Sommer et al., 2015; Broderick et al., 2014; Erkosar et al., 2015) and in human infants (Renz et al., 2012). To evaluate if exposing HIOs to *E. coli* led to maturation at the epithelial interface, we evaluated the transcriptional events following microinjection of live *E. coli* into the HIO lumen. PBS-injected HIOs (controls) and HIOs co-cultured with *E. coli* were collected for transcriptional analysis after 24, 48 and 96 hours (**Figure 2**). At 24 h post-microinjection, a total of 2,018 genes were differentially expressed (adjusted-FDR < 0.05), and the total number of differentially expressed genes was further increased at 48 and 96 h post-microinjection relative to PBS-injected controls (**Figure 2A**). Principle component analysis demonstrated that global transcriptional activity in HIOs is significantly altered by exposure to *E. coli*, with the degree of transcriptional change relative to control HIOs increasing over time (**Figure 2B**).

**Figure 2:**
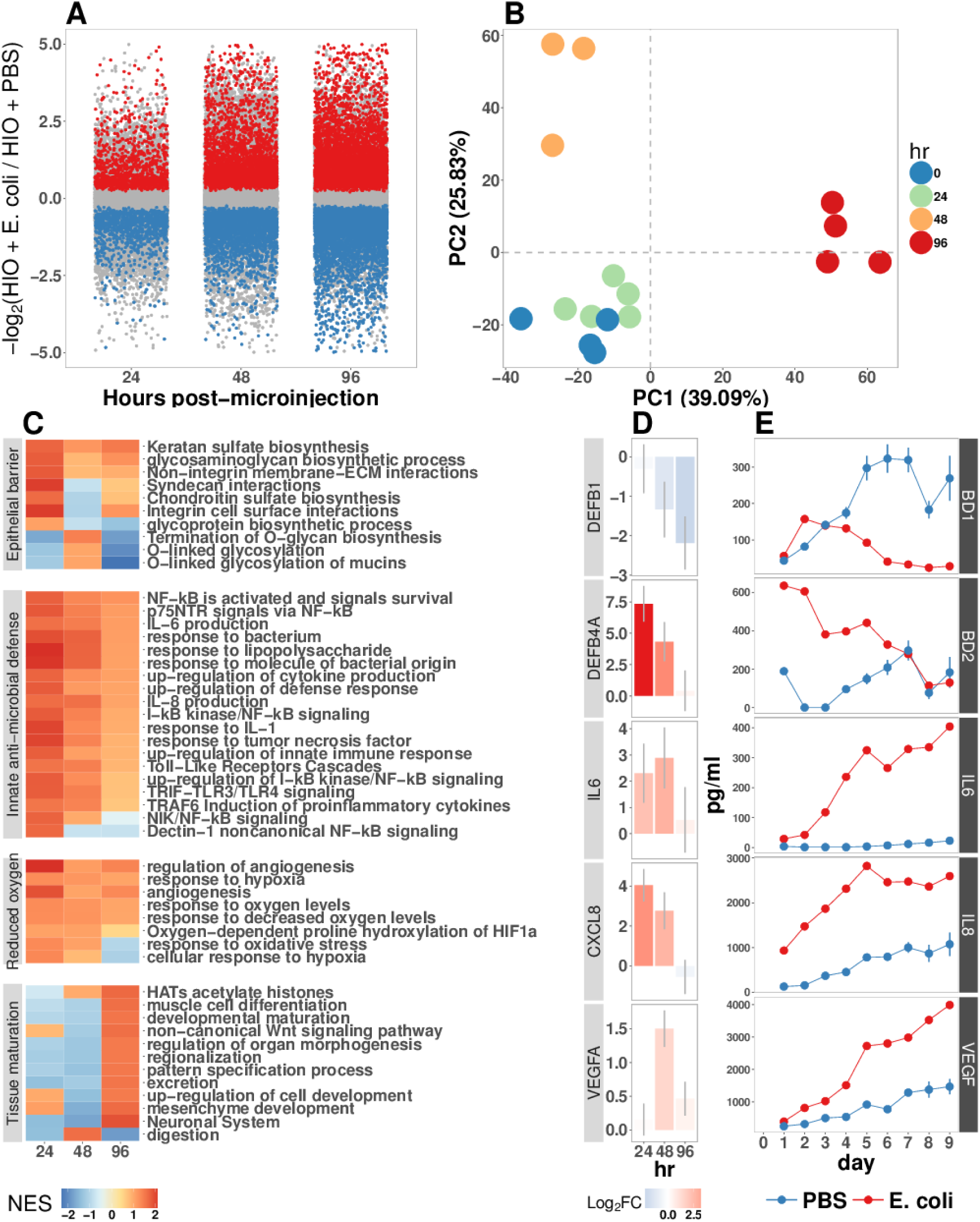
**A** Log2-transformed fold change in normalized RNA-seq gene counts in *E. coli* colonized HIOs at 24, 48, and 96 h post-microinjection with 10^4^ live *E. coli* relative to PBS-injected HIOs. Differentially expressed genes (FDR-adjusted *P*-value < 0.05) are indicated in red (up-regulated) or blue (down-regulated). Plotted results are the mean fold change per gene for each group. **B** Principle component plot of HIOs at 0-96 h post-microinjection derrived from whole-transcriptome RNA-seq normalized gene counts. Cumulative explained variance for PC1 and PC2 is indicated as a percentage on the x- and y-axes, respectively. **C** Heat map of normalized enrichment scores (NES) from GSEA of normalized RNA-seq expression data using the GO and REACTOME databases. A positive value of NES indicates activation of a given gene set and a negative value suggests relative suppression of a gene set. All NES scores are calculated relative to PBS-microinjected controls. **D** Mean log_2_ fold change in normalized RNA-seq gene counts at 24-96 h post microinjection relative to PBS-injected control HIOs. **E** Protein secretion at 0-9 days post-microinjection with PBS or *E. coli* as measured by ELISA in the supernatant of HIO cultures. The genes given in D correspond to the proteins measured in E. *N* = 4 (0 h), 5 (24 h), 3 (48 h), and 4 (96 h) biological replicates consisting of 5-6 pooled HIOs per replicate for panels A-D. *N* = 48 *E. coli*-injected HIOs and *N* = 8 PBS-injected HIOs for panel E.

Gene set enrichment analysis (GSEA) (Subramanian et al., 2005) using the GO (Ashburner et al., 2000; Gene Ontology Consortium, 2015) and REACTOME (Croft et al., 2014; Fabregat et al., 2016) databases to evaluate RNA-seq expression data revealed coordinated changes in gene expression related to innate anti-microbial defense, epithelial barrier production, adaptation to low oxygen, and tissue maturation (**Figure 2C**). Innate anti-microbial defense pathways, including genes related to NF-*κ*B signaling, cytokine production, and Toll-like receptor (TLR) signaling were strongly up-regulated at 24 h post-microinjection and generally exhibited decreased activation at later time points. GSEA also revealed changes in gene expression consistent with reduced oxygen levels or hypoxia, including the induction of pro-angiogenesis signals. A number of pathways related to glycoprotein synthesis and modification, including O-linked mucins, glycosaminoglycans, and proteoglycans, were up-regulated in the initial stages of the transcriptional response (Syndecans, integrins), exhibited a somewhat delayed onset (O-linked mucins), or exhibited consistent activation at all time points post-microinjection (Keratan sulfate and glycosaminoglycan biosynthesis). Finally, genes sets associated with a range of processes involved in tissue maturation and development followed a distinct late-onset pattern of expression. This included broad gene ontology terms for organ morphogenesis, developmental maturation, and regionalization as well as more specific processes such as differentiation of mesenchymal and muscle cells, and processes associated with the nervous system (**Figure 2C**).

We also made correlations between upregulated genes in the RNA-seq data (**Figure 2D**) and protein factors present in the organoid culture media following *E. coli* microinjection (**Figure 2E**). *β*-defensin 1 (*DEFB1* (gene); BD-1 (protein)) and *β*-defensin 2 (*DEFB4A* (gene); BD-2 (protein)) exhibited distinct patterns of expression, with both *DEFB1* and its protein product BD-1 stable at 24 hours after *E. coli* microinjection but relatively suppressed at later time points, and *DEFB4A* and BD-2 strongly induced at early time points and subsiding over time relative to PBS-injected controls. By contrast, inflammatory regulators IL-6 and IL-8 and the pro-angiogenesis factor VEGF were strongly induced at the transcriptional level within 24-48 h of *E. coli* microinjection. Secretion of IL-6, IL-8, and VEGF increased over time, peaking at 5 - 9 days after *E. coli* association relative to PBS-injected controls (**Figure 2E**). Taken together, this data demonstrates a broad-scale and time-dependent transcriptional response to *E. coli* association with distinct early- and late-phase patterns of gene expression and protein secretion.

### Bacterial colonization results in a transient increase in epithelial proliferation and the maturation of enterocytes

While the transcriptional analysis demonstrated strong time-dependent changes in the cells that comprise the HIO following E. coli colonization, we hypothesized that exposure to bacteria may also alter the cellular behavior and/or composition of the HIO. Previous studies have demonstrated that bacterial colonization promotes epithelial proliferation in model organisms (Bates et al., 2006; Cheesman et al., 2011; Neal et al., 2013; Kremer et al., 2013; Ijssennagger et al., 2015). We examined epithelial proliferation in HIOs over a timecourse of 96 hours by treating HIOs with a single 2 h exposure of 10 *μ*M EdU added to the culture media from 22-24 h after microinjection with 10^4^ CFU *E. coli* or PBS alone. HIOs were subsequently collected for immunohistochemistry at 24, 48, and 96 h post-microinjection (**Figure 3**). The number of proliferating epithelial cells (Edu^\+^ and E-cadherin^\+^) was elevated by as much as 3-fold in *E. coli*-colonized HIOs relative to PBS-treated HIOs at 24 h (**Figures 3A-B**). However, at 48 h post-microinjection, the proportion of EdU+ epithelial cells was significantly decreased in *E. coli* colonized HIOs relative to control treated HIOs. This observation was supported by another proliferation marker, KI67 (Gerdes et al., 1984)(**Figure 3B**), as well as RNA-seq data demonstrating an overall suppression of cell cycle genes in *E. coli* colonized HIOs relative to PBS-injected HIOs at 48 h post-microinjection (**Figure 3 - Supplement 1**). By 96 h post-microinjection the proportion of EdU+ epithelial cells was nearly identical in *E. coli* and PBS-treated HIOs (**Figure 3B**). Collectively, these results suggest that *E. coli* colonization is associated with a rapid burst of epithelial proliferation, but that relatively few of the resulting daughter cells are retained subsequently within the epithelium.

**Figure 3:**
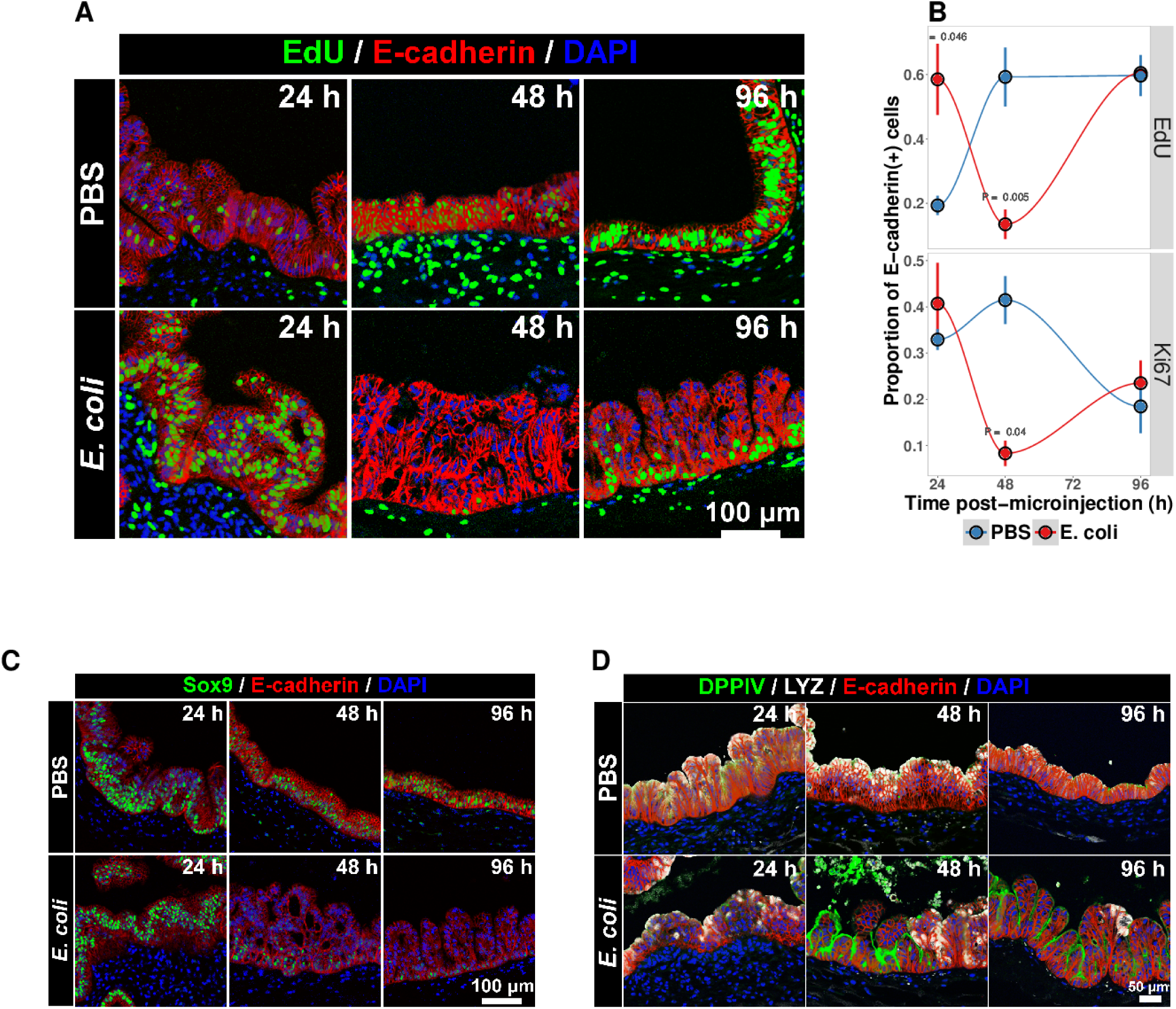
**A** Representative confocal micrographs of HIOs injected with PBS or 10^4^ CFU *E. coli* str. ECOR2 at 24-96 h post-microinjection and stained with fluorescent indicators for for EdU^+^ DNA, E-cadherin, or nuclei (DAPI) as indicated in the Figure labels. All HIOs were exposed to 10 *μ*M EdU at 22 h post-microinjection and EdU was removed at 24 h. Panels are representative of 4 HIOs per timepoint per treatment condition. **B** Quantification of the number of EdU-positive and Ki67-positive epithelial cells (E-cadherin^+^ cells) per 10X confocal microscopy field. One 10X confocal microscopy field consisting of 200-1000 epithelial cells was collected from each of 4 HIOs per timepoint per treatment group. The error bars represent the standard error of the mean and the *P*-values reflect the results of an unpaired two-tailed Student’s *t*-test comparing the PBS-injected HIOs to the *E. coli*-injected HIOs at that timepoint. **C** Representative confocal micrographs of HIOs injected with PBS or 10^4^ CFU *E. coli* str. ECOR2 at 24-96 h post-microinjection and stained with fluorescent antibodies for Sox9, E-cadherin, or nuclei (DAPI) as indicated in the Figure labels. Panels are representative of 4 HIOs per timepoint per treatment condition.

The transcription factor Sox9 is expressed by progenitor cells in the intestinal epithelium (Bastide et al., 2007; ***Mori-Akiyama et al., 2007***) and several epithelial subtypes are derived from a Sox9-expressing progenitor population in the mature intestinal epithelium (Bastide et al., 2007; Furuyama et al., 2011). We examined SOX9 expression in HIOs following microinjection with *E. coli* or PBS alone over a 96 hour time course (**Figure 3C**). In the PBS-treated HIOs, the majority of epithelial cells exhibited robust nuclear SOX9 expression at all time points examined. However, SOX9 expression was dramatically reduced in *E. coli*-colonized HIOs at 48-96 h after microinjection and was notably distributed in nuclei farthest from the lumen and adjacent to the underlying mesenchyme, mirroring the altered distribution of EdU+ nuclei seen in **Figure 3B**. This observation suggests that there is a reduction in the number of progenitor cells in the HIO epithelium following *E. coli* colonization and implies that other epithelial types may account for a greater proportion of the HIO epithelium at later time points post-colonization. We saw no appreciable staining for epithelial cells expressing goblet, Paneth, or enteroendocrine cell markers (MUC2, DEFA5, and CHGA, respectively; negative data not shown). However, expression of the small intestinal brush border enzyme dipeptidyl peptidase-4 (DPPIV) was found to be robustly expressed in the *E. coli*-colonized HIOs at 48 and 96 h post-microinjection (**Figure 3D**). DPPIV was not detected in any of the PBS-injected HIOs at any timepoint. Lysozyme (LYZ), an antimicrobial enzyme expressed by Paneth-like progenitorsin the small intestinal crypts ***Bevins and Salzman*** (***2011***), was widely distributed throughout the epithelium of PBS-treated HIOs as we have previously described (Spence et al., 2011)(**Figure 3D**). However, in *E. coli*-colonized HIOs, LYZ expression was restricted to distinct clusters of epithelial cells and, notably, never overlapped with DPPIV staining (**Figure 3D**). Given that *bonafide* Paneth Cell markers (i.e. DEFA5) were not observed in any HIOs, it is likely that the LYZ expression is marking a progenitor-like population of cells. Taken together, these experiments indicate that *E. coli* colonization induces a substantial but transient increase in the rate of epithelial proliferation followed by a reduction and redistribution of proliferating epithelial progenitors and differentiation of a population of cells expressing small intestinal enterocyte brush boarder enzymes over a period of 2-4 days.

### *E. coli* colonization is associated with a reduction in luminal O_2_

The mature intestinal epithelium is characterized by a steep oxygen gradient, ranging from 8% oxygen within the bowel wall to < 2% oxygen in the lumen of the small intestine (Fisher et al., 2013). Reduction of oxygen content in the intestinal lumen occurs during the immediate perinatal period (Gruette et al., 1965), resulting in changes in epithelial physiology (Glover et al., 2016; Kelly et al., 2015; Colgan et al., 2013; Zeitouni et al., 2016) that helps to shape the subsequent composition of the microbiota (Schmidt and Kao, 2014; Espey, 2013; Albenberg et al., 2014; Palmer et al., 2007; Koenig et al., 2011). Analysis of the global transcriptional response to *E. coli* association in the immature intestinal tissue revealed pronounced and coordinated changes in gene expression consistent with the onset of hypoxia (**Figure 2C-E**). We therefore measured oxygen concentration in the lumen of control HIOs and following microinjection of live *E. coli* using a 50 *μ*m diameter fiberoptic optode (**Figure 4A-B**). Baseline oxygen concentration in the organoid lumen was 8.04 ± 0.48%, which was significantly reduced relative to the external culture media (18.86 ± 0.37%, *P*= 3.6 × 10^−11^). At 24 and 48 h post-microinjection, luminal oxygen concentration was significantly reduced in *E. coli*-injected HIOs relative to PBS-injected HIOs (*P* = 0.04 and *P* = 5.2 × 10^−5^, respectively) reaching concentrations as low as 1.67 ± 0.62% at 48 h (**Figure 4A**). *E. coli* injected HIOs were collected and CFU were enumerated from luminal contents at 24 and 48 h post-microinjection. We observed a highly significant negative correlation between luminal CFU and luminal oxygen concentration where increased density of luminal bacteria was correlated with lower oxygen concentrations (r^2^ = 0.842, *P* = 6.86 × 10^−5^; **Figure 4B**). Finally, in order to assess relative oxygenation in the epithelium itself, we utilized a small molecule pimonidazole (PMDZ), which forms covalent conjugates with thiol groups on cytoplasmic proteins only under low-oxygen conditions (Arteel et al., 1998). Fluorescent immunochemistry demonstrated enhanced PMDZ uptake in *E. coli* associated HIO epithelium, and in HIOs grown in 1% O_2_ as a positive control when compared to to PBS-injected HIOs, or HIOs injected with heat killed *E. coli* at 48 h post-microinjection (**Figure 4C**). Thus, luminal and epithelial oxygen is reduced following microinjection of *E. coli* into the HIO, consistent with data in mice showing that the in *vivo* epithelium is in a similar low-oxygen state in normal physiological conditions (Schmidt and Kao, 2014; Kelly et al., 2015; Kim et al., 2017).

**Figure 4:**
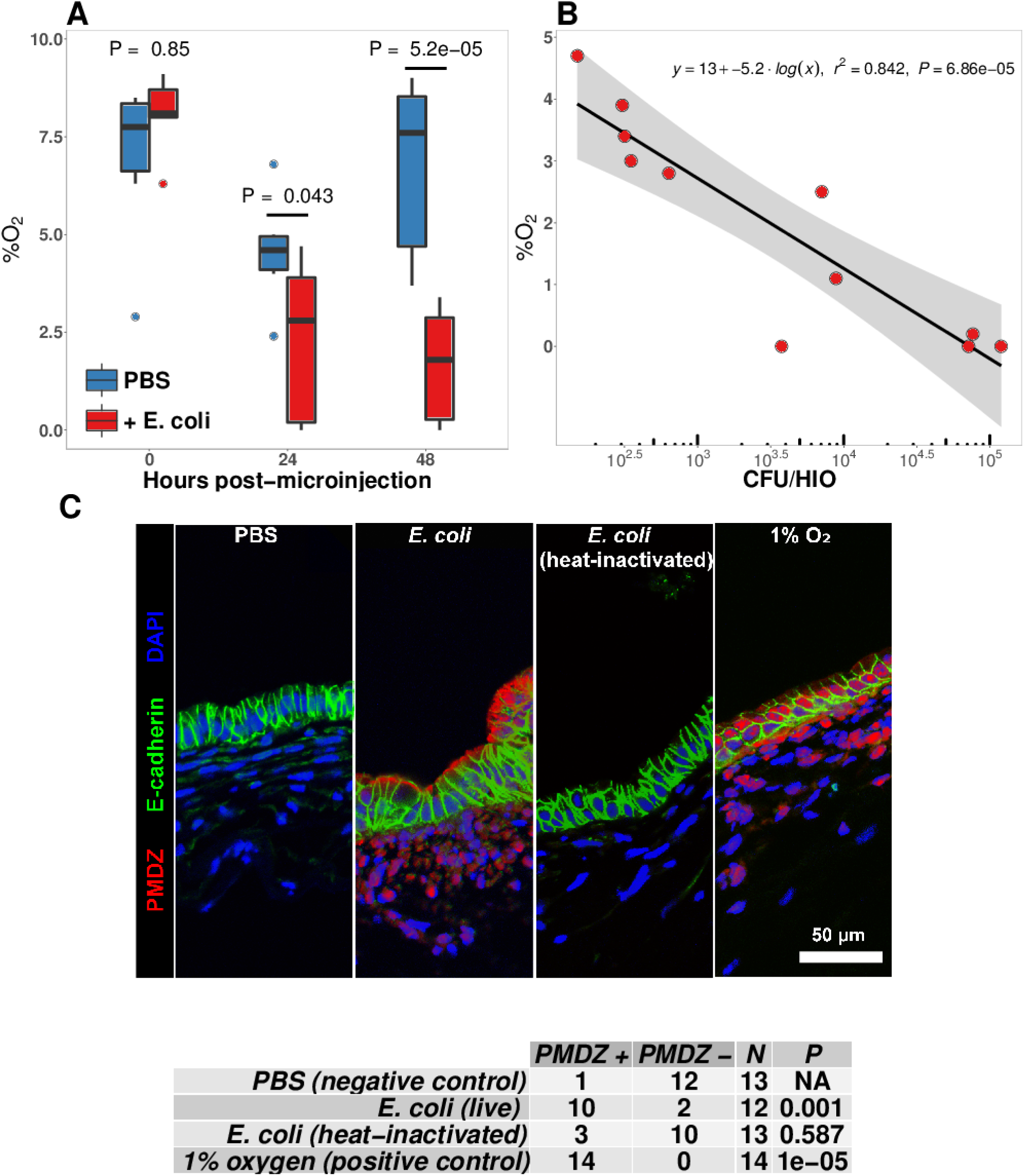
**A** Luminal oxygen concentration in human intestinal organoids at 0-48 h post-microinjection with 10^4^ CFU live *E. coli*. *P* values reflect results of unpaired one-tailed Students *t*-tests for the comparisons indicated. *N* = 6-11 replicate HIOs per treatment group per time point. **B** Linear regression analysis of luminal CFU *E. coli* per organoid at and luminal oxygen concentration in the same organoid 24 h post-microinjection. **C** Confocal micrographs of the HIO epithelium in PBS- and *E. coli*-injected HIOs at 48 h post-microinjection. Images are representative of the replicates detailed in the table, with 12-14 replicate HIOs per treatment group pooled from 2 separate experiments. Individual HIOs were scored as PMDZ^+^ or PMDZ^−^ based on the presence or absence, respectively, of PMDZ conjugates as detected by immunofluorescent microscopy. *P*-values represent the results of *x*^2^ contingency tests comparing the distribution of PMDZ^+^ and PMDZ^−^ HIOs in the PBS-treated group to each of the other conditions.

### NF-*к*B integrates complex microbial and hypoxic stimuli

*E. coli* colonization elicits a robust transcriptional response in immature intestinal tissue (**Figure 2**) that is associated with the onset of luminal oxygen depletion and relative tissue hypoxia (**Figure 4**). We set out to determine whether we could assign portions of the transcriptional response to direct interaction with microbes or to the subsequent depletion of luminal oxygen. In the RNA-seq analysis (**Figure 2**), NF-*к*B signaling emerged as a major pathway involved in this complex host-microbe interaction, and NF-*к*B has been shown by others to act as a transcriptional mediator of both microbial contact and the response to tissue hypoxia (Rius et al., 2008; Gilmore, 2006; Wullaert et al., 2011). Gene Ontology and REACTOME pathway analysis showed that NF-*к*B signaling components are also highly up-regulated following microinjection of *E. coli* into HIOs (**Figure 2C** and **Figure 5 - Supplement 1A**). Thus we assessed the role of NF-*к*B signaling in the microbial contact-associated transcriptional response and the hypoxia-associated response using the highly selective IKK*β* inhibitor SC-514 (Kishore et al., 2003; Litvak et al., 2009) to inhibit phosphorylation and activation of the transcription factor p65 (**Figure 5 - Supplement 1B**). Another set of HIOs was simultaneously transferred to a hypoxic chamber and cultured in 1% O_2_ with and without SC-514. At 24 h post-treatment, HIOs were harvested for RNA isolation and RNA-seq. We devised an experimental scheme that allowed us to parse out the relative contributions of microbial contact and microbe-associated luminal hypoxia in the transcriptional response to association with live *E. coli* (**Figure 5A** and **Figure 5 - Supplement 1C**). First, we identified a set of genes significantly up-regulated (log_2_FC > 0 & FDR-adjusted *P*-value < 0.05) by microinjection of either live *E. coli* or heat-inactivated *E. coli* (contact dependent genes). From this gene set, we identified a subset that was suppressed by the presence of NF-*к*B inhibitor SC-514 during association with either live or heat-inactivated *E. coli* (log_2_FC < 0 & FDR-adjusted *P*-value < 0.05; Gene Set I, **Figure 5B**). Thus, Gene Set I represents the NF-*к*B dependent transcriptional response to live or dead *E. coli*. Genes induced by live or heat-inactivated *E. coli* but not suppressed by SC-514 were considered NF-*к*B independent (Gene Set III, **Figure 5B**). Likewise, we compared genes commonly up-regulated by association with live *E. coli* and those up-regulated under 1% O_2_ culture conditions. A subset of genes induced by either live *E. coli* or 1% O_2_ culture but suppressed by the presence of NF-*к*B inhibitor was identified as the NF-*к*B-dependent hypoxia-associated transcriptional response (Gene Set II, **Figure 5B**). Genes induced by live *E. coli* or hypoxia but not inhibited by the presence of NF-*к*B inhibitor were considered NF-*к*B independent transcriptional responses to microbe-associated hypoxia (Gene Set IV). Gene lists for each gene set are found in **Supplementary File 1**. In **Figure 5C**, we examined the degree of overlap between Gene Set I, Gene Set II, and the set of genes that are significantly down-regulated in PBS-injected HIOs treated with SC-514. This analysis demonstrates that the majority of genes in Set I and Set II are not significantly down-regulated in PBS-injected HIOs treated with SC-514.

**Figure 5:**
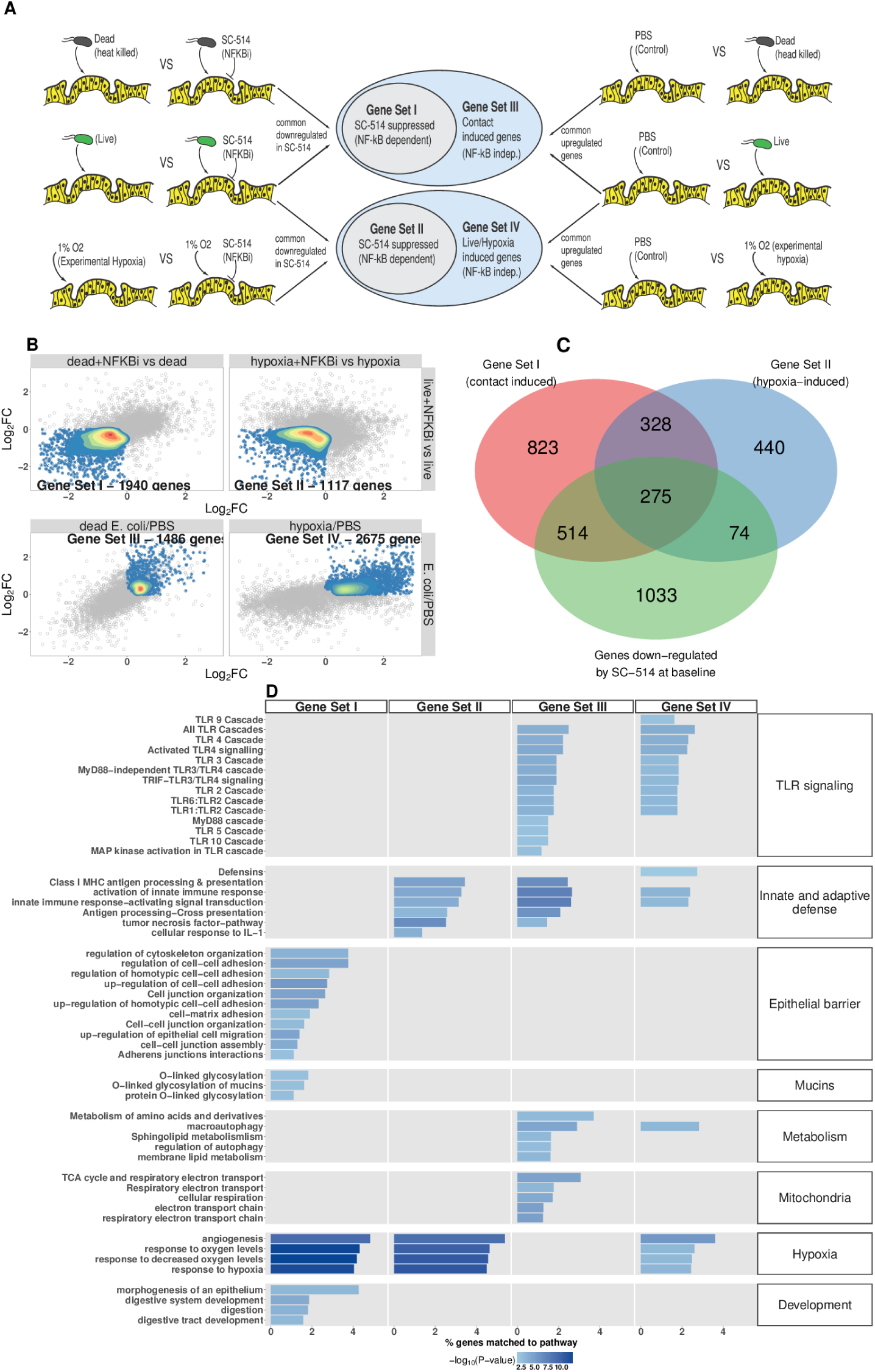
**A** Analysis scheme for identifying genes sets representing the components of the transcriptional response to live *E. coli* that could be recapitulated with heat-inactivated *E. coli* (contact induced) or hypoxia (microbial-associated hypoxia induced) as well as the subsets of genes induced through NF-*к*B dependent signaling. HIOs were microinjected with PBS, 10^4^ CFU *E. coli* or an equivalent concentration of heat-inactivated *E. coli* and cultured under standard cell culture conditions or hypoxic conditions (1% O_2_, 5% CO_2_, 94% N_2_) with and without 10 *μ*M SC-514. **B** Scatter plots with density overlay indicating the genes meeting the *a priori* criteria identified in panel A with an FDR-adjusted *P*-value of < 0.05 for the comparisons listed on the axes of the plot. **C** Venn Diagram showing the number of genes shared between Gene Set I, Gene Set II, and the set of genes that are significantly down-regulated in PBS-injected HIOs treated with SC-514 relative to PBS-injected control HIOs. **D** Bar plot of the proportion of genes in the input gene sets mapping to each pathway from the GO and REACTOME databases enrichment *P*-values for each of the gene sets identified in A. Pathways with enrichment *P*-values > 0.01 were excluded from the plot. Results represent *N* = 4-5 biological replicates per treatment condition, with each replicate consisting of 5-6 pooled and identically treated HIOs.

Following the identification of these 4 gene sets, we then applied over-representation analysis using the GO and REACTOME pathway databases to identify enriched pathways for each of the 4 gene sets, resulting in 4 clearly distinguishable patterns of gene pathway enrichment (**Figure 5D**). Contact with either live or heat-inactivated *E. coli* is sufficient to promote expression of genes involved in maintaining epithelial barrier integrity and mucin production, an effect that is suppressed in the presence of NF-*к*B inhibitor. Additionally, key developmental pathways including epithelial morphogenesis, digestive tract development, and expression of digestive enzymes appear to be driven primarily by bacterial association and are largely NF-*к*B dependent. Robust innate and adaptive defense requires both bacterial contact and hypoxia, with some genes associated with antigen processing and cytokine signaling being NF-*к*B dependent (Gene Set II) and others associated with NF-*к*B independent gene sets (Gene Sets III & IV). Genes associated with antimicrobial defensin peptides were enriched only in the hypoxia asociated, NF-*к*B independent gene set (Gene Set IV), suggesting that antimicrobial peptides are regulated by mechanisms that are distinct from other aspects of epithelial barrier integrity such as mucins and epithelial junctions (Gene Set I). TLR signaling components were is broadly enhanced by live *E. coli* and associated with both microbial contact and hypoxia were largely NF-*к*B independent (Gene Sets III & IV). There was a notable transcriptional signature suggesting metabolic and mitochondrial adaptation to bacteria that was independent of NF-*к*B and primarily driven by bacterial contact rather than hypoxia (Gene Set III).

To interrogate the transcriptional changes influenced by SC-514 exposure, we examined over-represented genes sets from the GO and REACTOME databases in genes that were significantly up- or down-regulated by treatment with SC-514 alone (**Figure 5 - Supplement 1C and D**). Notably, SC-514 alone does not appear to have a strong effect on the pathways identified in **Figure 5D** as key NF-*κ*B-dependent responses to bacterial contact and/or hypoxia. The most significant effects of SC-514 alone among Gene Set I and Gene Set II genes are related to metabolism, redox state, and ribosomal dynamics (**Figure 5 - Supplement 1F**). Thus, the effect of SC-514 alone cannot account for the NF-*κ*B-dependent changes in innate and adaptive defense, epithelial barrier integrity, angiogenesis and hypoxia signaling, or intestinal development following bacterial contact and/or hypoxia during colonization.

Finally, we also examined the role of microbial contact and hypoxia in colonization-induced changes in AMP, cytokine, and growth factor secretion using ELISA (**Figure 5 - Supplement 2**). Consistent with findings from the RNA-seq data, these results indicate that there are diverse responses to bacterial contact and hypoxia. We observed cases where cytokines were induced by either microbial contact or hypoxia alone (IL-6), other cases where hypoxia appeared to be the dominant stimuli (BD-1), and a third regulatory paradigm in which the response to live *E. coli* evidently results from the cumulative influence of bacterial contact and hypoxia (BD-2, IL-8, VEGF). Taken together, this analysis demonstrates that association of immature intestinal epithelium with live *E. coli* results in a complex interplay between microbial contact and microbe-associated hypoxia induced gene expression and protein secretion.

### Bacterial colonization promotes secretion of antimicrobial peptides

Antimicrobial peptides (AMPs) are key effectors for innate defense of epithelial surfaces (Muniz et al., 2012) and act to inhibit microbial growth through direct lysis of the bacterial cell wall and modulation of bacterial metabolism (Ganz, 2003; Bevins and Salzman, 2011; O’Neil, 2003; Vora et al., 2004; Brogden, 2005). Defensin gene expression is highly up-regulated following microinjection of *E. coli* into HIOs (**Figures 2D-E and 4C**). Using an annotated database of known AMPs (Wang et al., 2016) to query our RNA-seq datasets, we found that several AMPs are up-regulated in the immature intestinal epithelium following *E. coli* association (**Figure 6A**). Among these, DEFB4A and DEFB4B, duplicate genes encoding the peptide human *β*-defensin 2 (Harder et al., 1997), were the most highly up-regulated; other AMPs induced by *E. coli* association included multi-functional peptides CCL20, CXCL2, CXCL1, CXCL6, CXCL3, REG3A (Cash et al., 2006), and LTF (**Figure 6A**). Analysis of RNA-seq data from HIOs microinjected with live or heat-killed *E. coli* with and without NF-*к*B inhibitor or culture of HIOs under hypoxic conditions had indicated that defensin genes were enriched among the set of NF-*к*B-independent genes induced by hypoxia (**Figure 5D**). We examined *DEFB4A*expression specifically (**Figure 6B**) and found that relative to control treatment, microinjection of live *E. coli* resulted in a 7.38-fold increase in normalized *DEFB4A* expression. Consistent with the notion that *DEFB4A* expression is induced by hypoxia and is not dependent on NF-*к*B signaling, NF-*к*B inhibitor treated HIOs injected with E. coli still showed an ~8-fold increase in gene expression and hypoxia-cultured HIOs showed a ~5.5 fold induction (**Figure 6B**). On the other hand, microinjection with heat-inactivated *E. coli* resulted in *DEFB4A* induction that was significantly lower relative to microinjection with live *E. coli* (*P* = 0.007. A similar pattern of expression was observed for *DEFB4B* (**Figure 6 - Supplement 1**).

**Figure 6:**
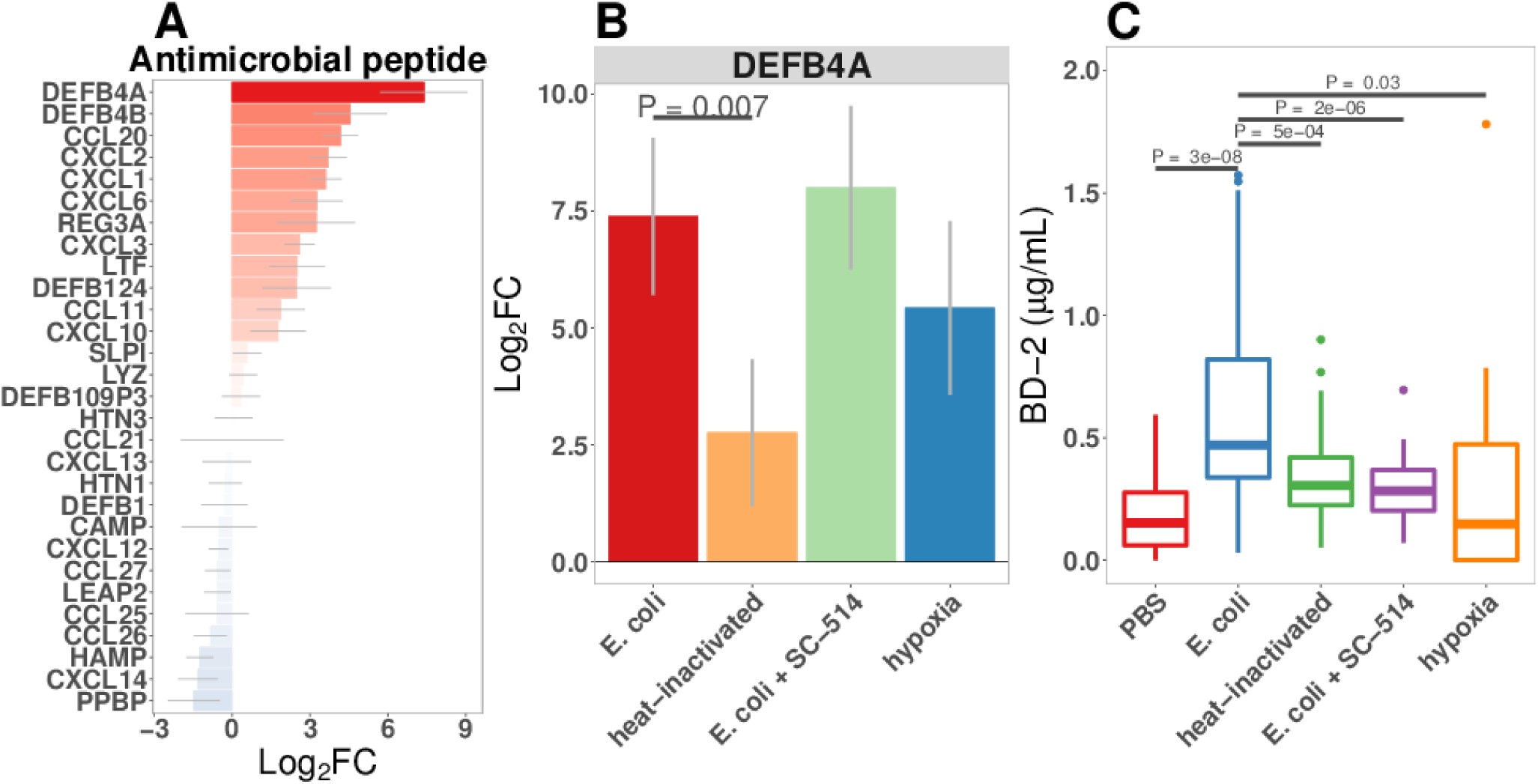
**A** Normalized fold change in antimicrobial peptide (AMP) gene expression in *E. coli*-associated HIOs at 24 h relative to PBS control treatment. **B** Normalized fold change in expression of DEFB4A, the gene encoding human *β*-defensin 2 (BD-2) peptide, in each of the conditions indicated relative to PBS control treatment. Results in panels A and B represent *N* = 4-5 biological replicates per treatment condition, with each replicate consisting of 5-6 pooled and identically treated HIOs. **C** Concentration of BD-2 peptide in culture supernatant at 24 h as measured by ELISA in HIO cultures treated as indicated. *N* = 10-14 individually treated HIOs per treatment condition with data combined from 3 independant replicate experiments. *P* values represents the results of a two-tailed Student’s *t*-test for the comparisons indicated.

We also examined secretion of human *β*-defensin 2 peptide (BD-2) in the supernatant of *E. coli*associated HIOs (**Figure 2E** and **Figure 6C**). BD-2 secretion was increased 3.4-fold at 24 h following *E. coli* microinjection (*P* = 2.7 × 10^−8^). However, heat-inactivation of *E. coli* or addition of NF-*к*B inhibitor resulted in suppression of BD-2 secretion relative to live *E. coli* (*P* = 0.000 51 and 1.6 × 10^−6^, respectively).

To determine if the levels of BD-2 produced by HIOs and secreted into the media were sufficient to retard bacterial growth, we tested the effect of BD-2 at concentrations consistent those found in HIO-/E. coli/ supernatant (1*μ*g/mL) (**Figure 6 - Supplement 2**). *E. coli* density was decreased over time in bacterial growth media supplemented BD-2, resulting in a significant decrease in the effective *in vitro* carrying capacity, or maximum population density (**Figure 6 - Supplement 2B**, *P* = 0.006). Additional data suggests that the inhibitory activity of BD-2 *in vitro* is not specific to *E. coli* str. ECOR2 and is dependent upon maintenance of BD-2 protein structure, since BD-2 similarly inhibited growth of *E. coli* str. K12, and heat-inactivated BD-2 lost these inhibitory effects (**Figure 6 - Supplement 2**). From this set of experiments we conclude that *E. coli* colonization promotes enhanced expression of AMPs, including BD-2, at concentrations that are sufficient to suppress microbial growth.

### Bacterial colonization promotes expression of epithelial mucins and glycotransferases

Mucins are an essential component of epithelial integrity, serving as a formidable barrier to microbial invasion and repository for secreted AMPs (Bergstrom and Xia, 2013; Cornick et al., 2015; Johansson and Hansson, 2016; Kim and Ho, 2010). Mucin synthesis requires a complex series of post-translational modifications that add high molecular weight carbohydrate side chains to the core mucin protein (Varki, 2017). Our RNA-seq data suggested that mucin gene expression is dependent on both bacterial contact and NF-*κ*B signaling (**Figure 5D**). Therefore, we examined expression of genes in control and *E. coli* microinjected HIOs that encode mucin core proteins as well as the glycotransferases that generate the wide variety of post-translational mucin modifications (**Figure 7A**). Although some glycotransferases were increased at 24 h after *E. coli* microinjection, expression of mucin core proteins and many glycotransferases reached peak levels at 48 h after the introduction of *E. coli* to the HIO lumen (**Figure 7A**). Periodic Acid-Schiff and Alcian blue staining (PAS/AB) of sections taken from HIOs at 48 h after *E. coli* microinjection reveal the formation of a robust mucin layer at the apical epithelial surface consisting of both acidic (AB-positive) and neutral (PAS-positive) glycoprotein components, suggesting a rich matrix of O-linked mucins, glycosaminoglycans, and proteoglycans (**Figures 7B-C**). Interestingly, we observed that *E. coli* association caused an initial induction of *MUC5AC* at 48h that was reduced by 96 h (**Figure 7A**). *MUC5AC* is most highly expressed within the gastric mucosa, but has also been reported in the duodenal epithelium (Buisine et al., 1998, ***2001***; Rodriguez-Pineiro et al., 2013). On the other hand, *MUC2* is more commonly associated with the duodenum, and increased more slowly, showing peak expression after 96 hours of association with *E. coli* (**Figure 7A**). Co-staining of control HIOs and *E. coli* microinjected HIOs demonstrated colocalization with *Ulex europaeus* agglutinin I (UEA1), a lectin with high specificity for the terminal fucose moiety Fuc*α*1-2Gal-R (**Figure 7D**). This suggests that following *E. coli* association, HIOs produce mucins with carbohydrate modifications associated with bacterial colonization *in vivo* (Cash et al., 2006; Hooper et al., 1999; Goto et al., 2014).

**Figure 7:**
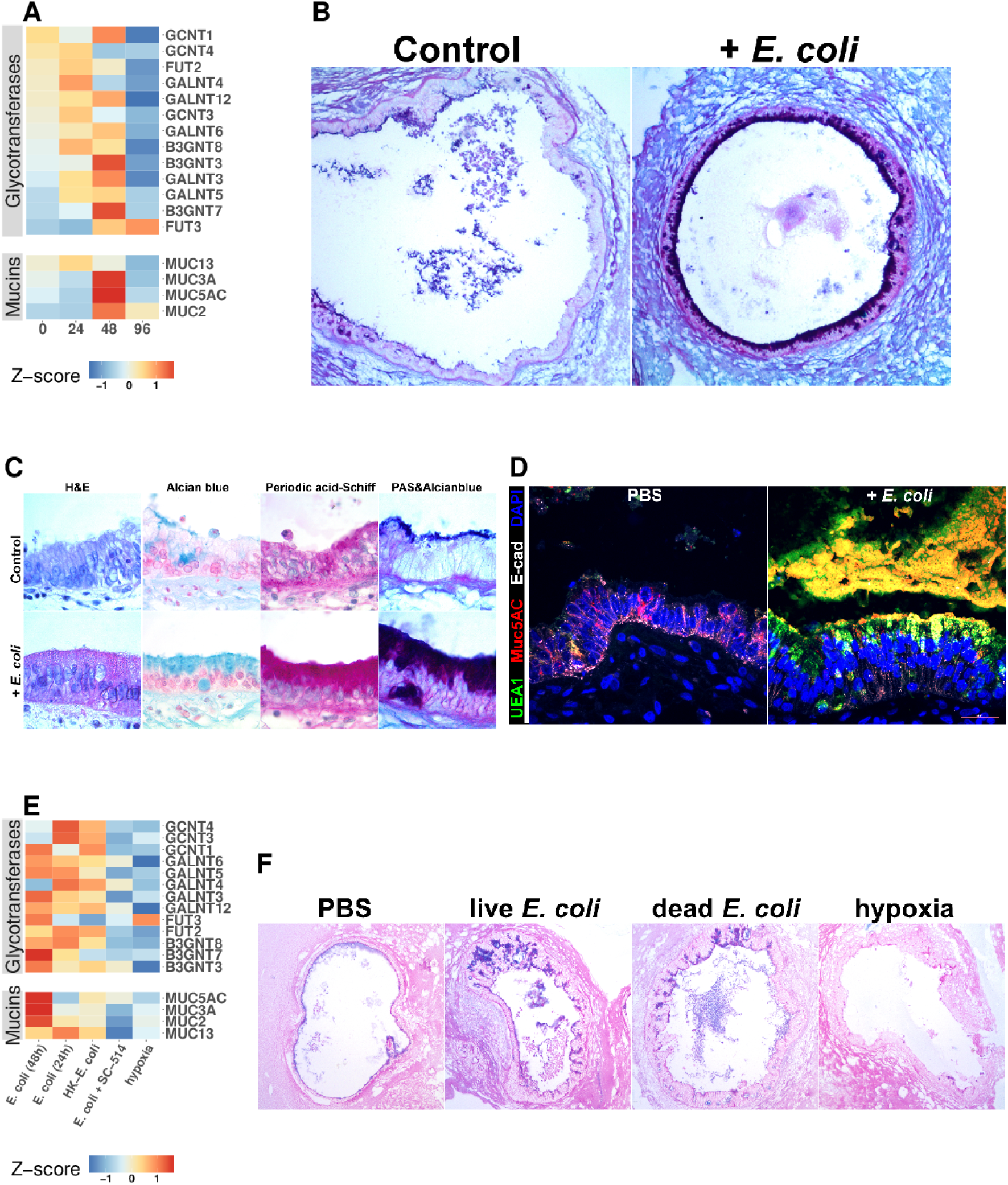
**A** Heatmap of normalized RNA-seq glycotransferase and mucin gene counts of HIOs associated with *E. coli* at 0-96 h post-microinjection. *N* = 4 (0 h), 5 (24 h), 3 (48 h), and 4 (96 h) biological replicates consisting of 5-6 pooled HIOs per replicate. **B** Periodic acid-Schiff and Alcian Blue (PAS-AB) staining of control HIOs or HIOs microinjected with *E. coli* and cultured for 48 h at 10X magnification. **C** HIO epithelium from control HIOs or HIOs microinjected with *E. coli* and cultured for 48 h stained with H&E, AB, PAS, or PAS-AB and imaged under 100X light microscopy. **D** Confocal micrograph of HIO epithelium from a control HIO or an HIO microinjected with *E. coli* and cultured for 48 h. Nuclei are stained blue with DAPI, and fluorescent antibody-labeled proteins E-cadherein and Mucin 5 AC are pseudocolored in white or red, respectively. UEA1 lectin is used to label the carbohydrate moiety Fuc*α*1-2Gal-R, which is pseudo colored in green. 60X optical magnification. **E** Heatmap of normalized RNA-seq glycotransferase and mucin gene counts of HIOs associated with live or heat-inactivated *E. coli*, *E. coli* + NF-*к*B inhibitor (SC-514) or HIOs cultured under hypoxic conditions for 24 h. Results represent the mean of *N* = 4-5 biological replicates per treatment condition, with each replicate consisting of 5-6 pooled and identically treated HIOs. **F** PAS-AB staining of HIOs treated as indicated in the figure labels for 24 h. 10X magnification. Histological and immunofluorescent images in panels B-D and F are representative of 3 or more independent experiments, each consisting of 5-10 HIOs per treatment group.

RNA-seq data suggested that O-linked mucins were highly enriched among the subset of genes induced by bacterial contact in an NF-*к*B-dependent manner (**Figure 5**). We examined this phenomenon at the level of individual glycosyltransferase and mucin genes (**Figure 7E**). *E. coli* induced transcription of mucins and glycosyltransferases (**Figure 7E**) and mucin secretion (**Figure 7 - Supplement 1**) was suppressed in the presence of NF-*к*B inhibitor SC-514. Furthermore, culture of HIOs under hypoxia conditions was not sufficient to promote transcription of genes involved in mucin synthesis (**Figure 7E**). This result was confirmed with PAS/AB staining of HIOs microinjected with PBS, live or heat-inactivated *E. coli*, or cultured under hypoxic conditions for 24 h, where bacterial contact promoted formation of a mucus layer while PBS microinjection or culture under hypoxic conditions did not (**Figure 7F**). Taken together, these results indicate that association of the immature intestinal epithelium with *E. coli* promotes robust mucus secretion through an NF-*к*B-dependent mechanism and that hypoxia alone is not sufficient to recapitulate *E. coli* induced mucus production.

### NF-*κ*B signaling is required for the maintenance of barrier integrity following bacterial colonization

Having established that the immature intestinal epithelium in HIOs (**Figure 1 - Supplement 1**) can be stably associated with non-pathogenic *E. coli* (**Figure 1**), resulting in broad changes in transcriptional activity (**Figure 2**) and leading to elevated production of AMPs (**Figure 6**) and epithelial mucus secretion (**Figure 7**), we hypothesized that these changes in gene and protein expression would have functional consequences for the immature epithelial barrier. RNA-seq analysis demonstrated broad up-regulation of transcription in genes involved in the formation of the adherens junction and other cell-cell interactions in HIOs after microinjection with live *E. coli* that was inhibited in the presence of NF-*к*B inhibitor SC-514 (**Figure 8A**). We utilized a modified FITC-dextran permeability assay (Leslie et al., 2015) and real-time imaging of live HIO cultures to measure epithelial barrier function in HIOs microinjected with PBS, live *E. coli*, or live *E. coli* + SC-514 at 24 h after microinjection (**Figure 8B**). While HIOs microinjected with PBS or *E. coli* retained 94.1 ± 0.3% of the FITC-dextran fluorescence over the 20 h assay period, *E. coli* microinjected HIOs cultured in the presence of SC-514 retained only 45.5 ± 26.3% of the fluorescent signal (*P* = 0.02; **Figure 8B**). We also measured the rate of bacterial translocation across the HIO epithelium, which resulted in contaminated culture media (**Figure 8C**). HIOs microinjected with *E. coli* and treated with SC-514 were compared to *E. coli* microinjected HIOs treated with vehicle (DMSO controls) and PBS microinjected controls over 7 days in culture. HIOs associated with *E. coli* + SC-514 exhibited a rapid onset of bacterial translocation by day 2-3, with bacterial translocation detected in 96% of SC-514 treated HIOs by day 7 compared to 23% of HIOs microinjected with *E. coli* and cultured in DMSO (*P* = *<*2 × 10^−16^; **Figure 8C**). Therefore, blocking NF-*к*B signaling inhibited epithelial barrier maturation resulting in increased bacterial translocation during *E. coli* association with the immature epithelium.

**Figure 8:**
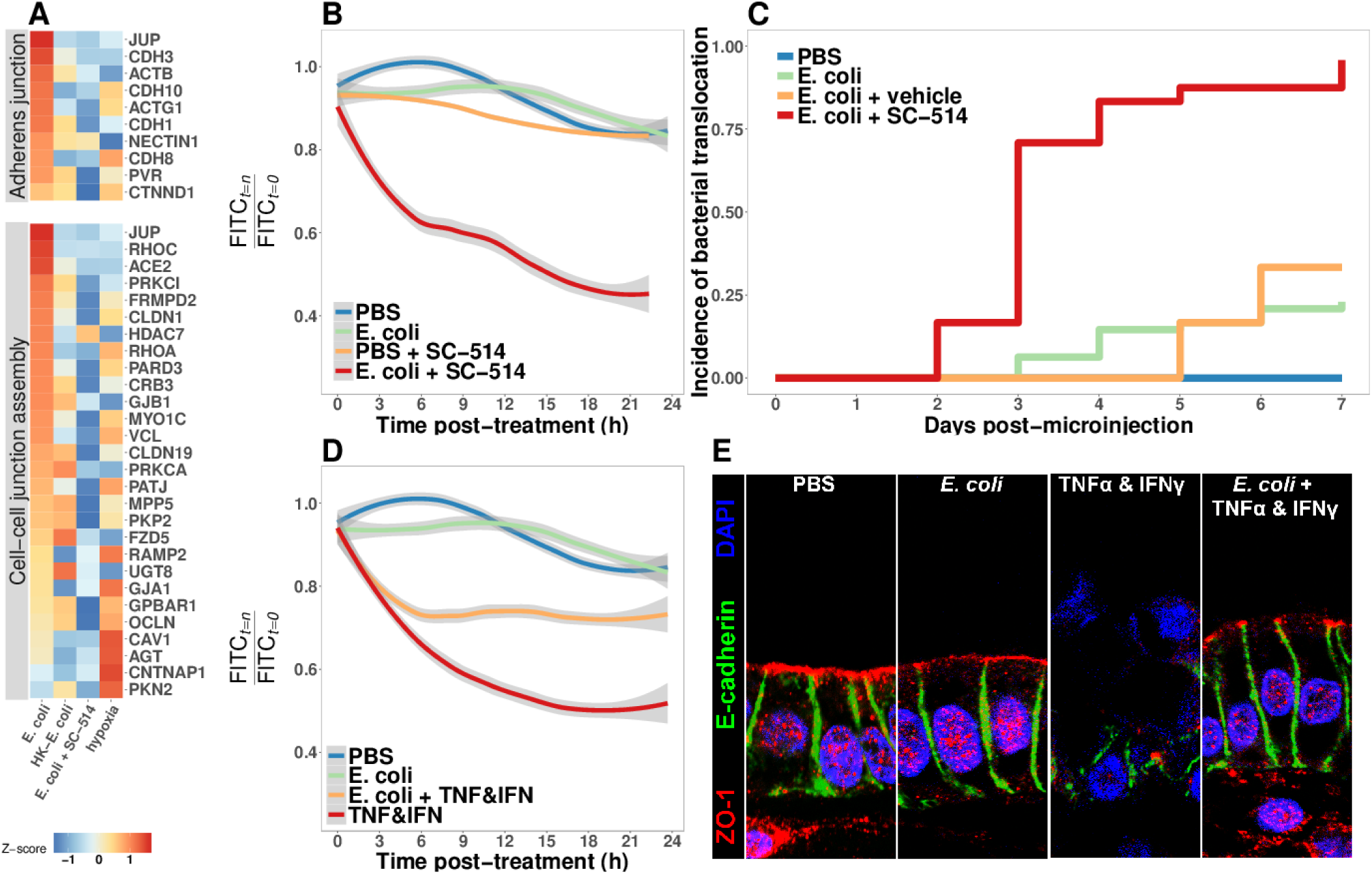
**A** Heatmap of RNA-seq data indicating the relative expression of genes associated with the Adherens junction or Cell-cell junction assembly based on annotation in the REACTOME database. Results represent the mean of *N* = 4-5 biological replicates per treatment condition, with each replicate consisting of 5-6 pooled and identically treated HIOs. **B** Relative fluoresscence intensity over time in HIOs microinjected with 4 kDa FITC-dextran and imaged at 10 minute intervals. HIOs were pretreated by microinjection with 10^4^ CFU *E. coli* in PBS or PBS alone and cultured for 24 h prior to treatment with media containing 10 *μ*M SC-514 or PBS alone and the injection of 2 mg/ml FITC-dextran (4 kDa) at the start of imaging. Line represents the best fit to the mean fluorescent intensity values in each condition with the grey region indicating S.E. for the fit line. *N*= 7-9 HIOs per group. **C** Rate of bacterial translocation over time in HIOs treated as indicated in the figure legend as detected by daily collection of external HIO media and enrichment in bacterial growth broth. *N* = 24 (*E. coli* + SC-514), *N* = 48 (*E. coli*), and *N* = 12 (PBS and *E. coli* + vehicle). **D** Relative fluorescence intensity over time in HIOs microinjected with FITC-dextran and imaged at 10 minute intervals. HIOs were pretreated by microinjection with 10^4^ CFU *E. coli* in PBS or PBS alone and cultured for 24 h prior to treatment with media containing 500 ng/ml TNF-*a* and 500 ng/ml IFN-*y* or PBS alone and the injection of 2 mg/ml FITC-dextran (4 kDa) at the start of imaging. Line represents the best fit to the mean fluorescent intensity values in each condition with the grey region indicating S.E. for the fit line. *N* = 8-9 HIOs per group. **E** Representative confocal micrographs of HIOs treated as indicated in D. Fluorescent immunostaining pseudocoloring applied as indicated in the figure legend. 60X optical magnification with 2X digital zoom. SC-514, small molecule inhibitor of NF-*к*B; HK, heat-inactivated; TNF, tumor necrosis factor-*a*; IFN, interferon-*y*

### Bacterial colonization promotes resilience of the epithelial barrier during cytokine challenge

Finally, we assayed epithelial barrier function under circumstances recapitulating physiologic inflammation. TNF*α* and IFN*γ* are key cytokines mediating innate and adaptive immune cell activity in the gut (Turner, 2009) during bacterial infection (Rhee et al., 2005; Emami et al., 2012) and in necrotizing enterocolitis (Tan et al., 1993; Ford et al., 1996, ***1997***; Halpern et al., 2003; Upperman et al., 2005). The combination of TNF*α* and IFN*γ* has been previously demonstrated to induce barrier permeability in a dose-dependent manner in Transwell epithelial cultures (Wang et al., 2005, ***2006***). Thus, HIOs were microinjected with PBS or live *E. coli* and cultured for 24 h, and were subsequently microinjected with FITC-dextran and treated with PBS or a cocktail of TNF*α* and IFN*γ* added to the external media to expose the basolateral epithelium (**Figure 8D**). Loss of FITC-dextran fluorescence was observed using live-imaging and indicated that treatment with TNF*α* and IFN*γ* alone resulted in a rapid and sustained decrease in luminal fluorescence relative to PBS or *E. coli* injected HIOs (*P* = 5 × 10^−4^, **Figure 8D**). However, HIOs associated with *E. coli* prior to addition of the TNF*α* and IFN*γ* cocktail retained significantly more fluorescent signal relative to treatment with TNF*α* and IFN*γ* alone (*P* = 0.042, **Figure 8D**). We examined expression and distribution of the tight junction protein ZO-1, and the basal-lateral protein E-cadherin (ECAD) in histological sections taken from PBS and *E. coli*-associated HIOs subjected to TNF*α* and IFN*γ* treatment (**Figure 8E**). Compared to controls, the epithelial layer is highly disorganized in HIOs treated with TNF*α* and IFN*γ*, with cytoplasmic ZO-1 staining and disorganized ECAD. By contrast, HIOs associated with *E. coli* prior to TNF*α* and IFN*γ* treatment retain and organized columnar epithelium with robust apical ZO-1 and properly localized ECAD staining (**Figure 8E**). Similarly, proper localization of additional markers of epithelial barrier integrity occludin (OCLN) and acetylated-tubulin are retained in HIOs associated with *E. coli* during TNF*α* and IFN*γ* treatment relative to HIOs treated with TNF*α* and IFN*γ* alone (**Figure 8 - Supplement 1**).These results suggest that colonization of the immature epithelium with *E. coli* results in an epithelium that is more robust to challenge by potentially damaging inflammatory cytokines.

## Discussion

The work presented here demonstrates that HIOs represent a robust model system to study the initial interactions between the gastrointestinal epithelium and colonizing microbes that occurs in the immediate postnatal period. Microorganisms introduced into the digestive tract at birth establish an intimate and mutualistic relationship with the host over time (Costello et al., 2012; Palmer et al., 2007; Koenig et al., 2011; Bäckhed et al., 2015; Wopereis et al., 2014). However, the expansion of bacterial populations in the gut also presents a major challenge to intestinal homeostasis through the exposure to potentially inflammatory MAMPs (Tanner et al., 2015; Renz et al., 2012), consumption of tissue oxygen (Glover et al., 2016; Espey, 2013; Albenberg et al., 2014), digestion of the mucus barrier (Marcobal et al., 2013; Desai et al., 2016), and competition for metabolic substrates (Rivera-Chávez et al., 2016; Kaiko et al., 2016). The mature intestinal epithelium serves as a crucial barrier to microbes that inhabit the lumen and mucosal surfaces (Artis, 2008; Turner, 2009; Desai et al., 2016; Kelly et al., 2015; Cornick et al., 2015; Peterson and Artis, 2014; Hackam et al., 2013; Turner, 2009). The specific function of the epithelium in adapting to initial microbial colonization, independent of innate and adaptive immune systems, remains unclear due to the lack of appropriate model systems that recapitulate host-microbe mutualism. Clarifying the role of the epithelium in colonization of the digestive tract by microorganisms is essential to understanding the molecular basis of the stable host-microbe mutualism in the mature intestine.

To examine the establishment of host-microbe mutualism, we chose to examine the interaction between the immature epithelium of HIOs and a non-pathogenic strain of *E. coli*. Enterobacteriaceae, including *E. coli*, are abundant in the newborn gut (Palmer et al., 2007; Koenig et al., 2011; Bäckhed et al., 2015; Yassour et al., 2016). Several large-scale surveys of microbial composition have demonstrated that *E. coli* are among the most prevalent and abundant organisms in stool samples from newborns (Bäckhed et al., 2015; Koenig et al., 2011) and in meconium (Gosalbes et al., 2013). Non-pathogenic *E. coli* strains may represent ideal model organisms for examining the impact of bacterial colonization of the immature epithelium due to their prevalence in the neonatal population and relevance to natural colonization, extensive characterization, and ease of laboratory manipulation. Microinjection of non-pathogenic *E. coli* into the lumen of HIOs resulted in stable, long-term co-cultures (**Figure 1**). *E. coli* grows rapidly within the HIO lumen (**Figure 1**), reaching densities roughly comparable to populations found in the human small intestine (Donaldson et al., 2016) within 24 h. Furthermore, the HIO is able to sustain this internal microbial population for several days while retaining the integrity of the epithelial barrier (**Figure 1**). Implicit is this observation is the conclusion that immature epithelium, along with a loosely structured mesenchymal layer, is intrinsically capable of adapting to the challenges imposed by colonization with non-pathogenic gut bacteria.

To more closely examine these epithelial adaptations of microbial colonization, we performed transcriptional analysis of this response. HIOs colonized by *E. coli* exhibit widespread transcriptional activation of innate bacterial recognition pathways, including TLR signaling cascades and downstream mediators such as NF-*κ*B (**Figure 2**). The cellular composition of the HIO epithelium is refined following *E. coli* colonization, with a rapid but transient increase in epithelial proliferation preceding a general reduction in the number of immature epithelial progenitor cells and the emergence of mature enterocytes expressing brush border digestive enzymes (**Figure 3**). Together, these results suggest that bacterial stimuli exert a broad influence on the molecular and cellular composition of the immature epithelium.

Indirect stimuli resulting from microbial activity can also shape epithelial function (***Buffie and Pamer, 2013***), and the transcriptome of *E. coli*-colonized HIOs reflects a cellular response to reduced oxygen availability (**Figure 2**). Reduction of luminal O_2_ concentration occurs in the neonatal gut (Gruette et al., 1965; Fisher et al., 2013; Zheng et al., 2015), possibly as a result of the consumption of dissolved O_2_ by the anaerobic and facultative anaerobic bacteria that predominate in the intestinal microbiome in early life (Espey, 2013; Fanaro et al., 2003; Favier et al., 2002; Palmer et al., 2007), and the mature intestinal epithelium is hypoxic relative to the underlying lamina propria due to the close proximity to the anaerobic luminal contents (Glover et al., 2016; Kelly et al., 2015; Zheng et al., 2015). We measured luminal oxygen content and epithelial hypoxia in HIOs microinjected with live *E. coli*, finding that luminal oxygen concentration is reduced more than 10-fold relative to the surrounding media. This state of relative hypoxia extends into the epithelium itself and is correlated with increased microbial density (**Figure 4**). Thus, although HIOs lack the network of capillaries that play an essential role in tissue oxygen supply in the intestine, *E. coli*-colonized HIOs recapitulate *in vitro* the oxygen gradient present at the epithelial interface.

Colonization of the HIO by *E. coli* therefore comprises two broad stimuli: immediate exposure to contact-mediated signals such as MAMPs, and the onset of limiting luminal oxygen and epithelial hypoxia. Although the potential significance of exposure to microbial products in the context of tissue hypoxia is widely recognized in the setting of necrotizing enterocolitis (Tanner et al., 2015; Afrazi et al., 2014; Hackam et al., 2013; Neu and Walker, 2011; Upperman et al., 2005; Nanthakumar et al., 2011), this two factor signaling paradigm has not been well studied as a component of normal intestinal colonization and development. Using the HIO model system, it was possible to design experiments which separately examine the relative impact of microbial contact-mediated signals from microbe-associated hypoxic signals (**Figure 5** and **Figure 5 - Supplement 2**). This approach reveals that the full transcriptional response generated by the HIO following *E. coli* colonization is the product of both contact-dependent and hypoxia-dependent signals, with heat-inactivated *E. coli* or hypoxia alone recapitulating distinct subsets of the changes in gene expression observed in HIOs colonized with live *E. coli* (**Figure 5**). Future studies may examine the role of additional hypoxia-independent live microbe-associated stimuli, such as metabolic products (Kaiko et al., 2016) and viability-associated MAMPs (Sander et al., 2011), in mediating the epithelial response to initial bacterial colonization.

NF-*κ*B signaling has been implicated in the downstream response to both microbial contact-mediated signals (Zhang and Ghosh, 2001; Xiao and Ghosh, 2005; Kawai and Akira, 2007) and tissue hypoxia (Koong et al., 1994; Rius et al., 2008; Arias-Loste et al., 2015; Oliver et al., 2009; Zeitouni et al., 2016; Colgan et al., 2013; Grenz et al., 2012). Pharmacologic inhibition of NF-*к*B resulted in the suppression of both microbial contact- and hypoxia-associated gene expression in HIOs, inhibiting both contact-mediated epithelial barrier defense pathways and hypoxia-associated immune activation (**Figure 5**). NF-*к*B appears to play a key role in integrating the complex stimuli resulting from exposure to microbial products and the onset of localized hypoxia in the immature intestinal epithelium during bacterial colonization.

The molecular and cellular maturation of the intestine that occurs during infancy ultimately results in enhanced functional capacity (Lebenthal and Lebenthal, 1999; Sanderson and Walker, 2000; Neu, 2007). Bacterial colonization is associated with enhanced epithelial barrier function in gnotobiotic animals, including changes in the production of antimicrobial peptides and mucus (Vaishnava et al., 2008; Cash et al., 2006; Goto et al., 2014; García-Lafuente et al., 2001; Malago, 2015; Ménard et al., 2008). Defensins produced in the intestinal epithelium are critical mediators of the density and composition of microbial populations in the gut and protect the epithelium from microbial invasion (Kisich et al., 2001; Ostaff et al., 2013; Cullen et al., 2015; Salzman et al., 2003, ***2010***). Production of BD-2 is dramatically increased in HIOs immediately following *E. coli* colonization (**Figure 2, Figure 5 -Supplement 2 and Figure 6**), reaching concentrations that are sufficient to limit overgrowth of *E. coli* (**Figure 6** and **Figure 6 - Supplement 2**) without completely precluding potentially beneficial bacterial colonization (**Figure 1**). Secreted and cell-surface associated mucins form a physical barrier to microbes in the gut, act as local reservoirs of antimicrobial peptide, and serve as substrates for the growth of beneficial microorganisms (Desai et al., 2016; Johansson and Hansson, 2016; Cornick et al., 2015; Hansson, 2012; Li et al., 2015; Dupont et al., 2014; Bergstrom and Xia, 2013). The immature HIO epithelium produces a robust mucus layer consisting of both neutral and acidic oligosaccharides with terminal carbohydrate modifications following colonization with *E. coli* (**Figure 7**). Importantly, hypoxia alone does not result in the production of mucus while the introduction of heat-inactivated *E. coli* induces mucus secretion at the apical epithelium (**Figure 7**), suggesting that microbial contact is the major stimulus eliciting mucus secretion in HIOs.

Epithelial barrier permeability is an important parameter of intestinal function reflecting the degree of selectivity in the transfer of nutrients across the epithelial layer and the exclusion of bacteria and other potentially harmful materials (Bischoff et al., 2014). Increases in epithelial barrier permeability occur in the setting of inflammation (Ahmad et al., 2017; Michielan and D’Incà, 2015) and infectious disease (Shawki and McCole, 2017). Colonization of HIOs with *E. coli* results in increased transcription of genes associated with the formation of the adherens junction and other cell-cell interactions in the epithelium (**Figure 8**). However, inhibition of NF-*к*B signaling dramatically increases both epithelial barrier permeability and the rate of bacterial translocation (**Figure 8**), suggesting that NF-*к*B signaling is critical to maintaining epithelial barrier integrity following colonization. Expression of genes involved in the formation of the cell junction and the production of antimicrobial defensins and mucus are NF-*κ*B dependent (**Figures 6-8**, **Figure 7 - Supplement 1**, Tsutsumi-Ishii and Nagaoka 2002; Ahn et al. 2005). The inability to mount an effective innate defense response in the presence of NF-*к*B inhibition results in the failure of the HIO epithelial barrier and the loss of co-culture stability (**Figure 8**). This result underscores the critical role of NF-*к*B signaling in the formation of a stable host-microbe mutualism at the immature epithelial interface.

Dysregulated production of pro-inflammatory cytokines contributes to the loss of epithelial barrier integrity in NEC (Tanner et al., 2015; Hackam et al., 2013; Neu and Walker, 2011; Nanthakumar et al., 2011; Halpern et al., 2003; Ford et al., 1997, ***1996***; Tan et al., 1993); this is recapitulated in HIOs, as exposure to pro-inflammatory cytokines results in the rapid loss of epithelial barrier integrity and the dissolution of epithelial tight junctions (**Figure 8**). Probiotics may promote epithelial barrier integrity in NEC (Robinson, 2014; Alfaleh et al., 2011; Underwood et al., 2014; Khailova et al., 2009) and HIOs colonized by *E. coli* exhibit enhanced epithelial barrier resilience (**Figure 8**). Functional maturation resulting from colonization of the immature intestinal epithelium may therefore play an essential role in promoting the resolution of physiologic inflammation.

While great progress has been made in characterizing the composition of the gut microbiota in health and disease (Shreiner et al., 2015; Costello et al., 2012), this approach has a limited ability to discern the contributions of individual bacteria to the establishment of host-microbe symbiosis. Our work establishes an approach that recapitulates host-microbe mutualism in the immature human intestine in an experimentally tractable *in vitro* model system. Application of this approach may facilitate the development of mechanistic models of host-microbe interactions in human tissue in health and disease. For example, one of the major limitations in our understanding of NEC has been the lack of an appropriate model system to study colonization of the immature intestine (Neu and Walker, 2011; Balimane and Chong, 2005; Tanner et al., 2015; Nguyen et al., 2015). Our results suggest that colonization of the HIO with a non-pathogenic gut bacteria results in functional maturation of the epithelial barrier. Future work which examines the effects of organisms associated with the premature gut (Morrow et al., 2013; Greenwood et al., 2014; Ward et al., 2016) on the molecular, cellular, and functional maturation of the immature epithelium may be instrumental in elucidating mechanisms of microbiota-associated disease pathogenesis in the immature intestine.

## Materials and Methods

### HIO culture

Human ES cell line H9 (NIH registry #0062, RRID:CVCL_9773) was obtained from the WiCell Research Institute. H9 cells were authenticated using Short Tandem Repeat (STR) DNA profiling (Matsuo et al., 1999) at the University of Michigan DNA Sequencing Core and exhibited an STR profile identical to the STR characteristics published by ***Josephson et al.*** (***2006***). The H9 cell line was negative for *Mycoplasma* contamination. Stem cells were maintained on Matrigel (BD Biosciences, San Jose, CA) in mTeSR1 medium (STEMCELL Technologies, Vancouver, Canada). hESCs were passaged and differentiated into human intestinal organoid tissue as previously described (Spence et al., 2011; McCracken et al., 2011). HIOs were maintained in media containing EGF, Noggin, and R-spondin (ENR media, see ***McCracken et al.*** (***2011***)) in 50 *μ*l Matrigel (8 mg/ml) without antibiotics prior to microinjection experiments. For hypoxic culture experiments, HIOs were transferred to a hydrated and sealed Modular Incubator Chamber (MIC-101, Billups-Rothenburg, Inc. Del Mar CA) filled with 1% O_2_, 5% CO_2_, and balance N_2_ and maintained at 37 °C for 24 h.

### HIO transplantation and tissue derived enteroid culture

HIO transplantations: This study was performed in strict accordance with the recommendations in the Guide for the Care and Use of Laboratory Animals of the National Institutes of Health. All animal experiments were approved by the University of Michigan Institutional Animal Care and Use Committee (IACUC; protocol # PRO00006609). HIO transplants into the kidney capsule were performed as previously described (***Finkbeiner et al., 2015b***; Dye et al., 2016) Briefly, mice were anesthetized using 2% isofluorane. The left flank was sterilized using Chlorhexidine and isopropyl alcohol, and an incision was made to expose the kidney. HIOs were manually placed in a subcapsular pocket of the kidney of male 7–10 week old NOD-SCID ILflRgnull (NSG) mice using forceps. An intraperitoneal flush of Zosyn (100 mg/kg; Pfizer Inc.) was administered prior to closure in two layers. The mice were sacrificed and transplant retrieved after 10 weeks. Human Tissue: Normal, de-identified human fetal intestinal tissue was obtained from the University of Washington Laboratory of Developmental Biology. Normal, de-identified human adult intestinal tissue was obtained from deceased organ donors through the Gift of Life, Michigan. All human tissue used in this work was obtained from non-living donors, was de-identified and was conducted with approval from the University of Michigan IRB (protocol # HUM00093465 and HUM00105750). Isolation and culture of HIO epithelium, transplanted HIO epithelium, fetal and adult human duodenal epithelium was carried out as previously described (***Finkbeiner et al., 2015b***), and was cultured in a droplet of Matrigel using L-WRN conditioned medium to stimulate epithelial growth, as previously described (Miyoshi et al., 2012; Miyoshi and Stappenbeck, 2013)

### Bacterial culture

*Escherichia coli* strain ECOR2 (ATCC 35321) was cultured in Luria broth (LB) media or 1.5% LB agar plates at 37 °C under atmospheric oxygen conditions. Glycerol stock solutions are available upon request. The assembled and annotated genome for the isolate of *Escherichia coli* strain ECOR2 used in these studies is available at https://www.patricbrc.org/view/Genome/562.18521. *Esherichia coli* strain K-12 MG1655 (CGSC #6300) was obtained from the Coli Genetic Stock Center at Yale University (http://cgsc2.biology.yale.edu/) and was used only in the *in vitro* BD-2 activity experiments. Whole genome sequencing of *Escherichia coli* strain ECOR2 was performed by the University of Michigan Host Microbiome Initiative Laboratory using the Illumina MiSeq platform.

### Microinjection

Microinjections were performed using a protocol modified from ***Leslie et al.*** (***2015***). Briefly, HIOs were injected using thin wall glass capillaries (TW100F-4, World Precision Instruments, Sarasota FL) shaped using a P-30 micropipette puller (Sutter Instruments, Novato CA). Pulled microcapilaries were mounted on a Xenoworks micropipette holder with analog tubing (BR-MH2 & BR-AT, Sutter Instruments) attached to a 10 ml glass syringe filled with sterile mineral oil (Fisher Scientific, Hampton NH). Fine control of the micropippette was achieved using a micromanipulator (Narishge International Inc., East Meadow NY) and microinjection was completed under 1-2X magnification on an SX61 stereo dissecting scope (Olympus, Tokyo Japan). HIOs suspended in Matrigel (Corning Inc., Corning NY) were injected with approximately 1 *μ*l solution. A detailed and up-to-date HIO microinjection protocol is available at ***Hill*** (***2017b***). In bacterial microinjection experiments, the HIO culture media was removed immediately following microinjection and the cultures were rinsed with PBS and treated with ENR media containing penicillin and streptomycin to remove any bacteria introduced to the culture media during the microinjection process. After 1 h at 37 °C, the HIOs were washed again in PBS and the media was replaced with fresh anti-biotic free ENR.

### Measurement of luminal oxygen

Luminal oxygen content was measured in HIOs using an optically coated implantable microsensor with a tip tapered at < 50 *μ*m (IMP-PSt1, PreSens Precision Sensing GmbH) attached to a micro fiber optic oxygen meter (Microx TX3, PreSens Precision Sensing GmbH, Regensburg Germany). The oxygen probe was calibrated according to the manufacturer’s instructions and measurements of the external media and HIO luminal oxygen content were collected by mounting the microsensor on a micromanipulator (Narishge International Inc., East Meadow NY) and guiding the sensor tip into position using 1-2X magnification on a stereo dissecting scope (Olympus, Tokyo Japan). All oxygen concentration readings were analyzed using PreSens Oxygen Calculator software (TX3v531, PreSens Precision Sensing GmbH, Regensburg Germany). For measurement of relative cytoplasmic hypoxia, HIO cultures were treated with 100 *μ*M pimonidazol HCl (Hypoxyprobe, Inc., Burlington MA) added to the external culture media and incubated at 37 °C and 5% CO_2_ for 2 h prior to fixation in 4% parafomaldehyde. Pimonidazole conjugates were stained in tissue sections using the Hypoxyprobe-1 mouse IgG monoclonal antibody (Hypoxyprobe, Inc., Burlington MA, RRID:AB_2335667) with appropriate secondary antibody (see antibody dilutions table).

### Immunohistochemistry

Immunostaining was carried out as previously described (***Finkbeiner et al., 2015b***). Antibody information and dilutions can be found in **Supplementary File 3**. All images were taken on a Nikon A1 confocal microscope or an Olympus IX71 epifluorescent microscope. CarboFree blocking buffer (SP-5040; Vector Laboratories, Inc. Burlingame CA) was substituted for dilute donkey serum in PBS in staining for mucins and carbohydrate moieties. EdU treatment and EdU fluorescent labeling using Click-iT chemistry was applied according to the manufacturer’s instructions (#C10339 Thermo Fisher). **Supplementary File 2** contains a table of all primary and secondary antibodies, blocking conditions, and product ordering information.

### NF-*к*B inhibition

The NF-*κ*B inhibitor SC-514 (Kishore et al., 2003; Litvak et al., 2009) (Tocris Cookson, Bristol, United Kingdom) was re-suspended in DMSO at a concentration of 25 mM. HIOs were treated with SC-514 suspended in DMSO added to the external ENR culture media at a final concentration of 1 *μ*M. Efficacy of SC-514 was verified by Western blot of lysates from HIOs injected with PBS or live *E. coli* or injected with live *E. coli* in the presence of 1 *μ*M SC-514 added to the external media. HIOs were collected after 24 hours in lysis buffer composed of 300 mM NaCl, 50mM Tris base, 1mM EDTA, 10% glycerol, 0.5% NP-40, and 1X Halt Phosphatase Inhibitor Cocktail (Pierce Biotechnology, Rockford IL). Lysates were separated on a 10% Bis-Tris polyacrylamide gel under reducing conditions (Invitrogen, Carlsbad CA) and transferred to PVDF using a wet transfer apparatus (Bio-Rad Laboratories, Hercules CA) overnight at 4 °C. The PVDF membrane was blocked in Odyssey TBS blocking buffer (LI-COR Biosciences, Lincoln NE). The membrane was submerged in blocking buffer containing primary rabbit monoclonal antibodies against phosphorylated NF-*к*B p65 (1:200, Cell Signaling Technology #3033S) or total NF-*к*B p65 (1:400, Cell Signaling Technology #8242S) and incubated at room temperature for 2 h. All washes were conducted in Tris-buffered saline with 1% Tween-20 (TBST). The secondary goat anti-rabbit IgG IRDye 800CW was diluted 1:15,000 in TBST and exposed to the washed membrane for 1 h at room temperature. After additional washes, the PVDF membrane was imaged using an Odyssey Scanner (LI-COR Biosciences, Lincoln NE).

### Bacterial translocation assay

Incidence of bacterial translocation was determined in HIOs plated individually in single wells of 24-well plates and microinjected with *E. coli*. The external culture media was collected and replaced daily. The collected media was diluted 1:10 in LB broth in 96 well plates and cultured at 37 °C overnight. Optical density (600 nm) was measured in the 96-well LB broth cultures using a VersaMax microplate reader (Molecular Devices, LLC, Sunnyvale CA). OD_600_ > sterile LB broth baseline was considered a positive culture.

### FITC-dextran permeability

For epithelial permeability assays, HIOs were microinjected with 4 kDa FITC-dextran suspended in PBS at a concentration of 2 mg/ml as described previously (Leslie et al., 2015) using the microin-jection system detailed above. Images were collected at 10 minute intervals at 4X magnification on an Olympus IX71 epifluorescent microscope using a Deltavision RT live cell imaging system with Applied Precision softWoRx imaging software (GE Healthcare Bio-Sciences, Marlborough MA). Cultures were maintained at 37 °C and 5% CO_2_ throughout the imaging timecourse. For experiments involving cytokine treatment, recombinant TNF-*α* (#210-TA-010, R&D Systems) and INF-*γ* (#AF-300-02, Peprotech) were added to the external culture media at a concentration of 500 ng/ml at the start of the experiment. A detailed and up-to-date HIO microinjection and live imaging protocol is available at ***Hill*** (***2017b***).

### *In vitro* antimicrobial activity assay

Recombinant human BD-2 (Abcam, Cambridge MA) was reconstituted in sterile LB broth and diluted to 0.1-1 *μ*g/ml. *E. coli* cultures were diluted 1:1000 in sterile LB containing 0-1 *μ*/ml BD-2 and transferred to a 96-well microplate. A VersaMax microplate reader (Molecular Devices, LLC, Sunnyvale CA) was used to measure OD_600_ at 10 minute intervals in microplates maintained at 37 °C with regular shaking over a 18 h timecourse. For stationary phase anti-microbial assays, overnight cultures of *E. coli* str. ECOR2 were diluted in PBS containing 1 *μ*/ml BD-2 or heat-inactivated BD-2 (heated at 120 °C for > 60 min) and placed in a 37 °C bacterial incubator for 5 hours. Cultures were then spread on LB agar plates and cultured overnight. The number of CFU was counted manually.

### ELISA assays

Secreted cytokine, antimicrobial peptide, and growth factor concentrations were determined by ELISA (Duosets, R&D Systems, Minneapolis, MN) using the manufacturer’s recommended procedures at the Immunological Monitoring Core of the University of Michigan Cancer Center.

### RNA sequencing and analysis

RNA was isolated using the mirVana RNA isolation kit and following the "Total RNA" isolation protocol (Thermo-Fisher Scientific, Waltham MA). RNA library preparation and RNA-sequencing (single-end, 50 bp read length) were performed by the University of Michigan DNA Sequencing Core using the Illumina Hi-Seq 2500 platform. All sequences were deposited in the EMBL-EBI ArrayExpress database (RRID:SCR_004473) using Annotare 2.0 and are cataloged under the accession number **E-MTAB-5801**. Transcriptional quantitation analysis was conducted using 64-bit Debian Linux stable version 7.10 ("Wheezy"). Pseudoalignment of RNA-seq data was computed using kallisto v0.43.0 (Bray et al., 2016) and differential expression of pseudoaligned sequences was calculated using the R package DEseq2 (Love et al., 2014) (RRID:SCR_000154).

### Statistical analysis

Unless otherwise indicated in the figure legends, differences between experimental groups or conditions were evaluated using an unpaired Student’s *t*-test. A *P*-value < 0.05 was considered to represent a statistically significant result. All statistical analyses were conducted using R version 3.4.2 (2017-09-28) (R Core Team, 2017) and plots were generated using the R package ggplot2 (Wickham, 2009) with the ggstance expansion pack ***Henry et al.*** (***2016***). The multiple testing-adjusted FDR was calculated using the DESeq2 implementation of the Wald test (Love et al., 2014). Gene pathway over-representation tests and Gene Set Enrichment Analysis (Subramanian et al., 2005) were implemented using the R packages clusterProfiler (Yu et al., 2012) and ReactomePA (Yu and He, 2016). Analyses conducted in R were collated using Emacs v25.2 (Stallman, 1981) with Org-mode v8.3.5 and the paper was written in L^A^T_E_X using Emacs. Complete analysis scripts are available on the ***Hill*** (***2017a***) GitHub repository.

## Acknowledgments

The authors would like to thank Joel Whitfield of the Immunological Monitoring Core of the University of Michigan Cancer Center, Robert Lyons and Tricia Tamsen of the University of Michigan DNA Sequencing Core, Chris Edwards of the University of Michigan Molecular Imaging Laboratory, and April Cockburn and Micah Kiedan of the University of Michigan Host Microbiome Initiative Laboratory for providing invaluable technical assistance. JRS is supported by the Intestinal Stem Cell Consortium (U01DK103141), a collaborative research project funded by the National Institute of Diabetes and Digestive and Kidney Diseases (NIDDK) and the National Institute of Allergy and Infectious Diseases (NIAID). JRS and VBY are supported by the NIAID Novel Alternative Model Systems for Enteric Diseases (NAMSED) consortium (U19AI116482). DRH is supported the Mechanisms of Microbial Pathogenesis training grant from the National Institute of Allergy and Infectious Disease (NIAID, T32AI007528) and the National Center for Advancing Translational Sciences (ULFITR000433).

## Supplementary File Legends

**Video 1** Animation of individual epifluorescent microscopy images from a human intestinal organoid (HIO) containing live GFP^+^ *E. coli* str. ECOR2. Images were captured at 10 minute intervals over the course of 18 h an coalated in sequential order. Representative of 3 independent experiments. See Figure 1A.

**Supplementary File 1** List of differentially expressed genes and the Gene Set (I-IV) assignments used in the analyses presented in Figure 5.

**Supplementary File 2** Immunostaining conditions for all immunohistochemistry data presented in this manuscript. NDS, Normal donkey serum; PBS, phosphate buffered saline.

---

**Figure 1 - Supplement 1.**
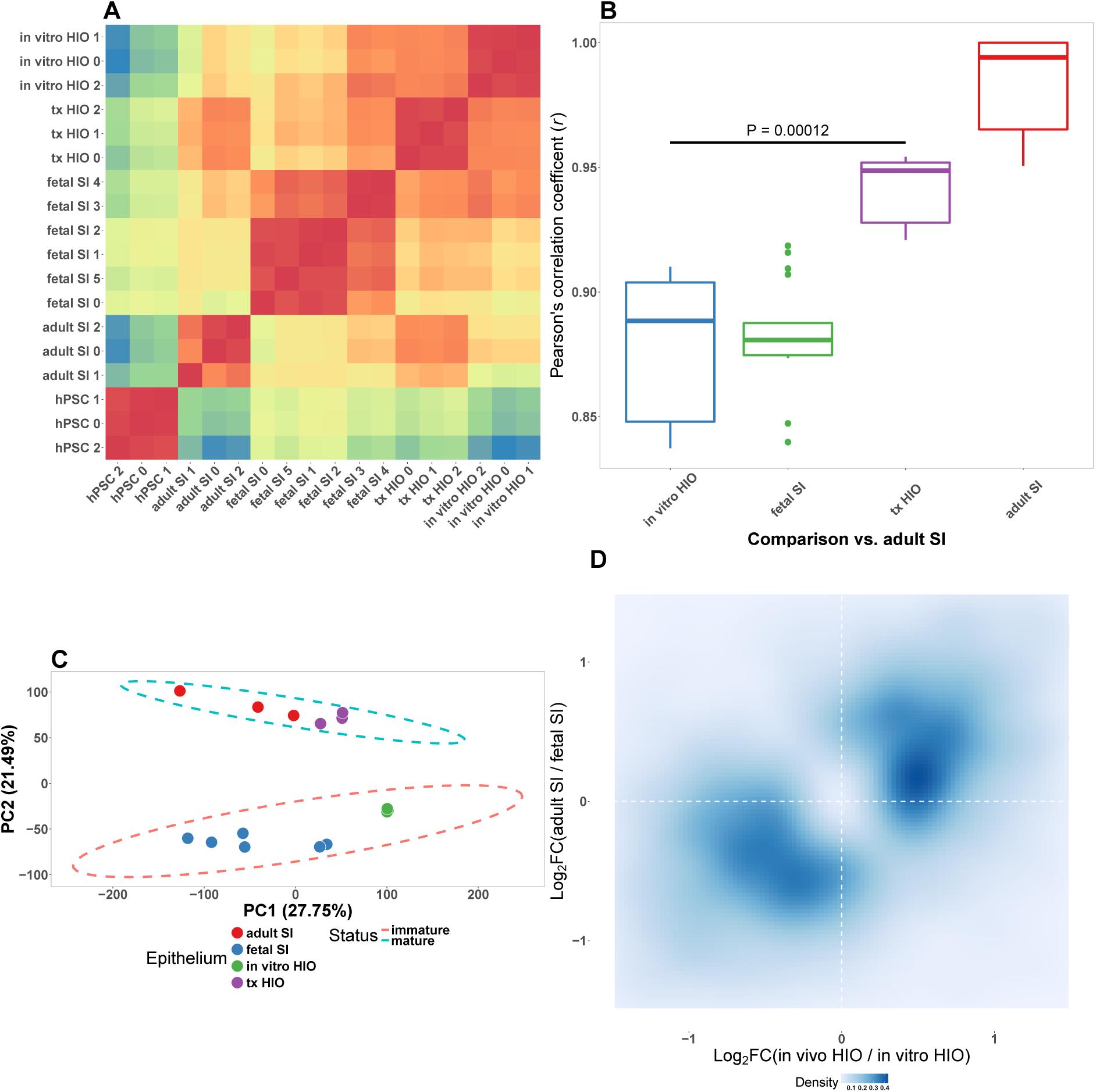
**A** Pearson’s correlation matrix with heirarchical clustering for whole-transcriptome normalized RNA-seq gene counts from epithelium isolated from the tissues indicated on the axes. **B** Pearson’s correlation coefficient for the comparison of whole-transcriptome normalized RNA-seq gene counts between each of the sample types listed on the x-axis and adult small intestinal epithelium. *P*-value indicates the results of an unpaired two-sided Student’s *t*-test. **C** Principle component analysis of whole-transcriptome RNA-seq normalized gene counts. Cumulative explained variance for PC1 and PC2 is indicated as a percentage on the x- and y-axes, respectively. **D** Density plot of the Log_2_-transformed Fold change in gene expression in epithelium from transplanted HIOs over epithelium from HIOs cultured *in vitro* plotted against the Log_2_-transformed Fold change in gene expression in adult small intestinal epithelium over fetal small intestinal epithelium. The intensity of the blue color indicates the density of points in 2-dimensional space. SI, small intestine; tx, transplanted tissue; hPSC, human pluripotent stem cell; HIO, human intestinal organoid

**Figure 1 - Supplement 2.**
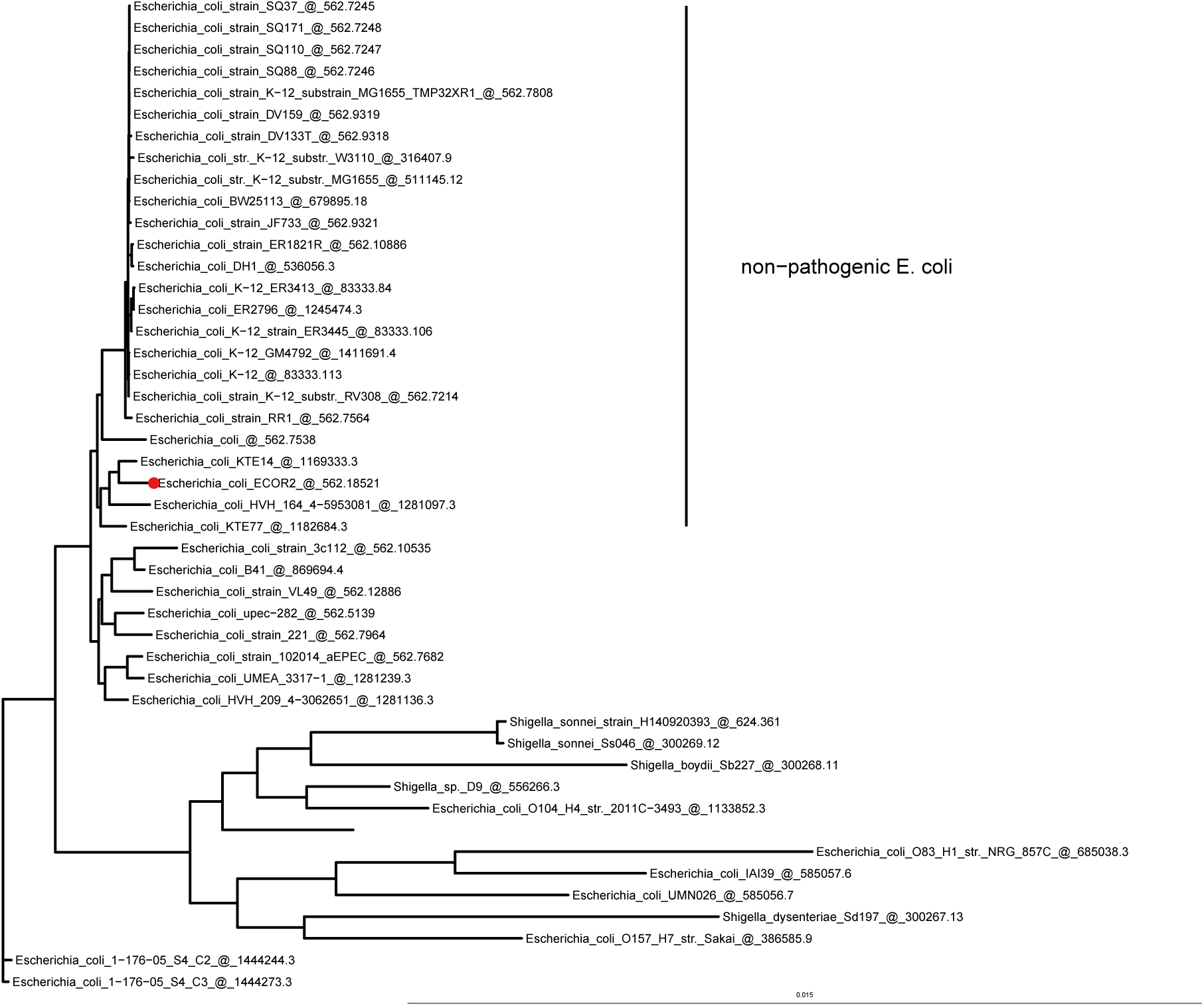
Phylogenetic tree based on maximum liklihood genomic distance among *E. coli* str. ECOR2 (***Ochman and Selander, 1984***), the strain used in the HIO colonization experiments, closely related *E. coli* isolates available on the PATRIC (***Wattam et al., 2017***) database, and pathogenic type strains from the genera *Esherichia*, *Shigella*, and *Salmonella*. A PATRIC genome reference number follows name of each taxa.

**Figure 1 - Supplement 3.**
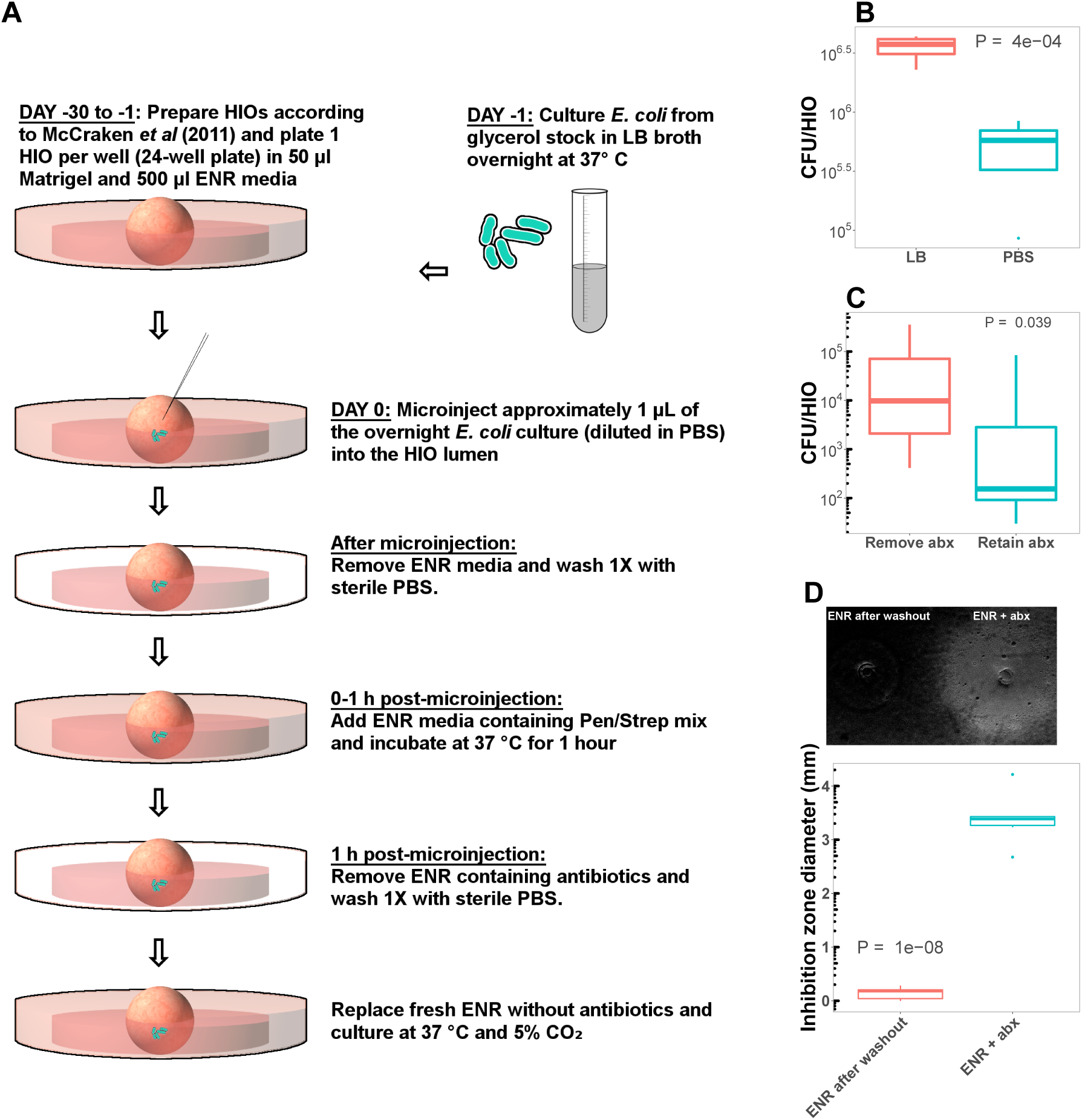
**A** Schematic representation of the microinjection of HIOs with live *E. coli*. See Materials and Methods for additional details. **B** A comparison of CFU/HIO at 24 h post-microinjection of HIOs microinjected with 10 × 10^3^ CFU live *E. coli* diluted in sterile PBS or fresh LB broth. *N* = 5 HIOs per condition. All experiments presented in the main paper represent *E. coli* diluted in PBS. *C* To test for the effect of antibiotic carryover of *E. coli* growth in the HIO lumen, we compared CFU/HIO in HIOs cultured in antibiotic free media ("Remove abx") or media containing penicillin and streptomycin at 24 h post-microinjection with 10^3^ CFU live *E. coli*. All experiments presented in the main paper represent HIOs cultured in antibiotic-free media due to the apparent effect of antibiotics in suppressing *E. coli* growth within the HIO lumen. *N* = 5 HIOs per condition. *D* In several experiments, bacterial translocation was measured by samplign the external HIO culture media (Figures 1 and 8). To evaluate the potential influence of antibiotic carryover in our estimates of bacterial translocation, we measured growth inhibition in *E. coli* cultures plated as a lawn of LB agar and treated with 1 *μ*l samples of HIO media collected during the 1 h antibiotic wash step or after HIO culture washout with PBS and replacement with fresh antibiotic free media (see panel A). *N* = 6 replicates per culture condition. All *P*-values represent the results of unpaired one-tailed Student’s *t*-test comparisons.

**Figure 3 - Supplement 1.**
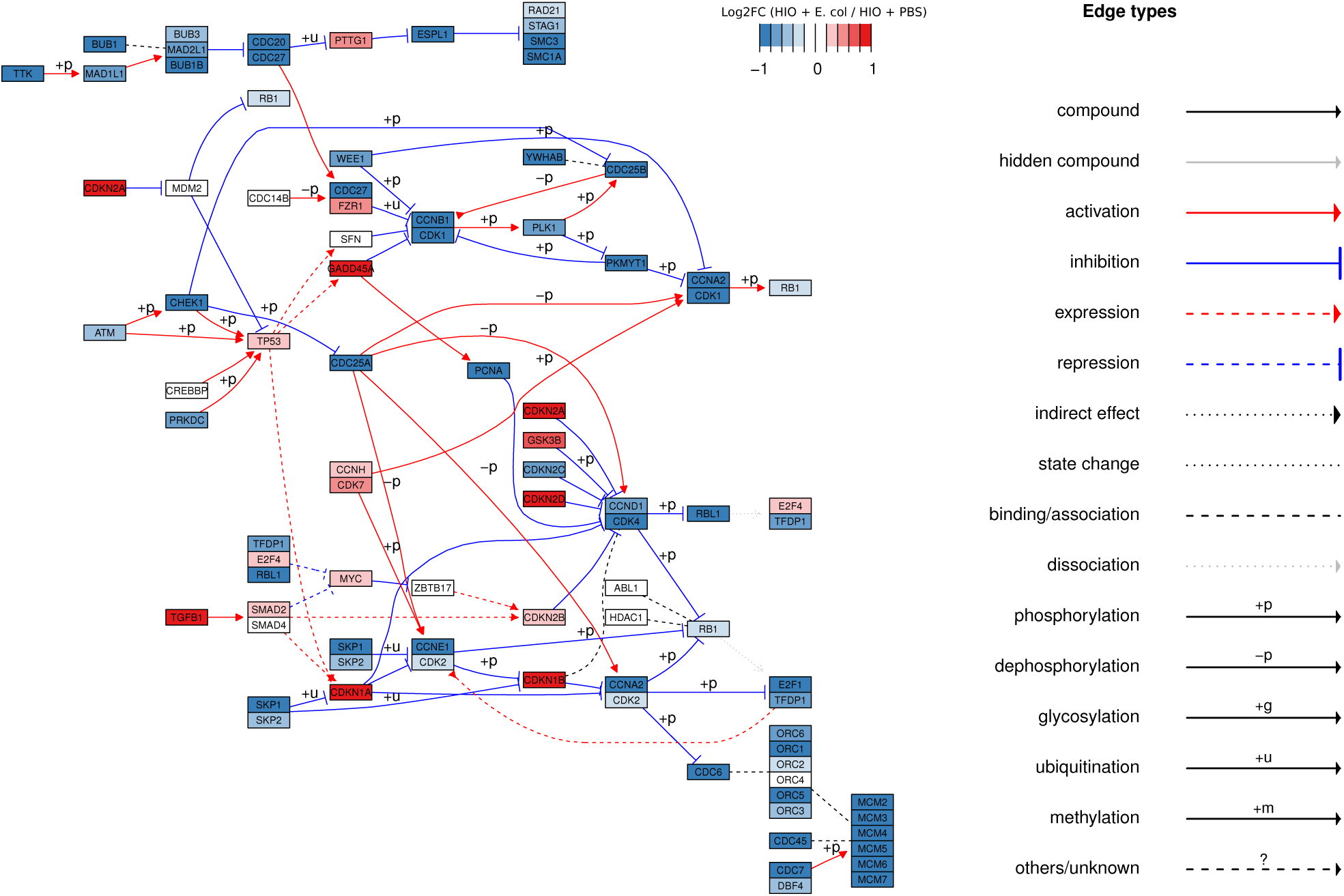
Pathview (**?**) plot of the KEGG (**?**) pathway showing the Cell mitotic cycle (“HSA 04110”) superimposed with RNA-seq expression data corresponding to the log_2_-transformed fold change in expression of cell cycle transcripts from HIOs microinjected with *E. coli* relative to PBS-injected HIOs at 48 h post-microinjection. Mean of 4 biological replicates, each representing 5-6 pooled HIOs.

**Figure 5 - Supplement 1.**
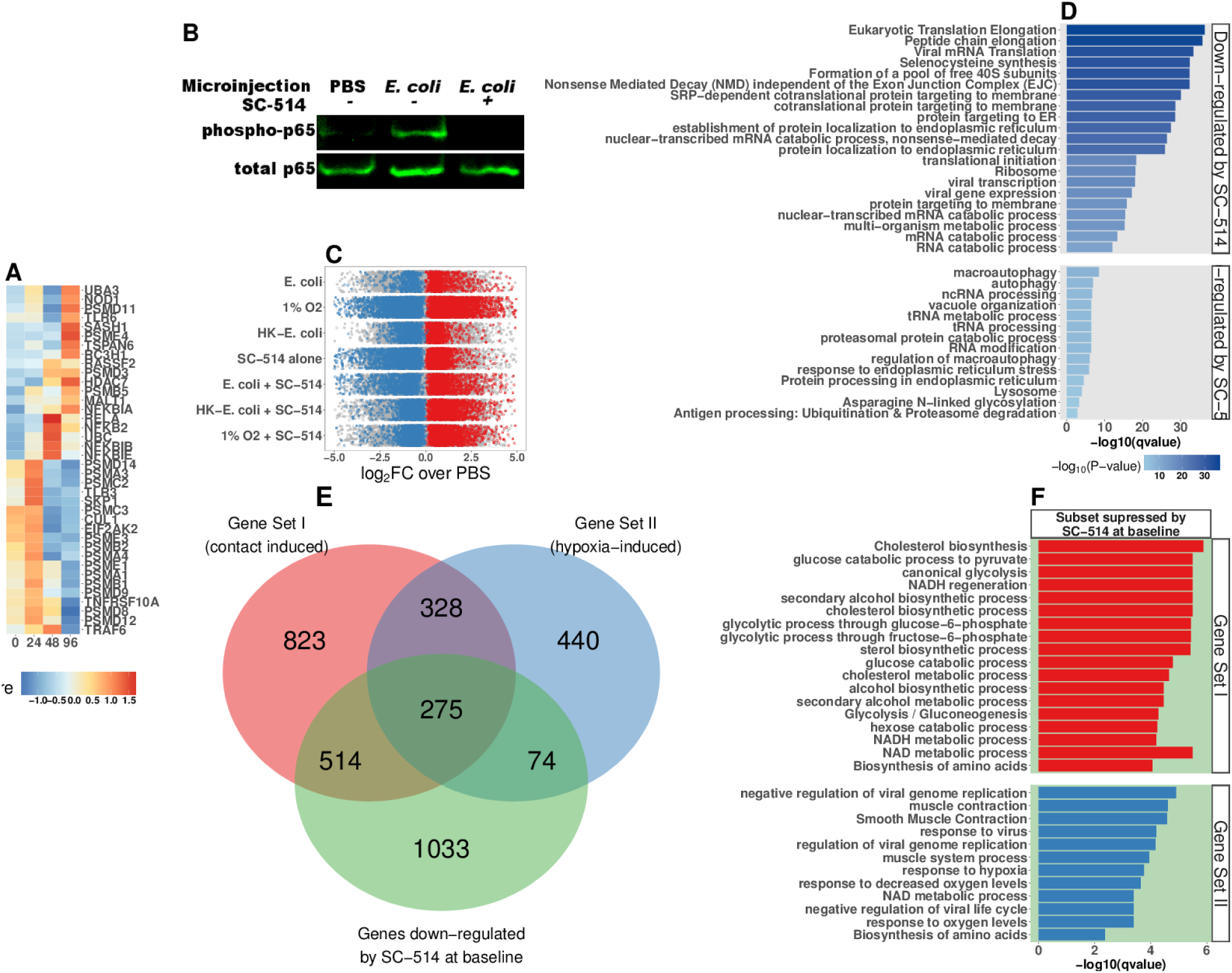
**A** Heatmap representation of time-dependent normalized RNA-seq gene expression for NF-*к*B pathway components in HIOs at 0-96 h post-microinjection. **B** Western blot of phosphorylated p65 and total p65 in cell lysates from HIOs microinjected with PBS or live *E. coli* and treated with IKK*/1* inhibitor SC-514 (1 *μ*M) as indicated in the figure. **C** log2-transformed fold-change in gene expression relative to PBS-injected controls for all 7 experimental conditions examined in this set of experiments: live E. coli +/- SC-514, heat-killed E. coli +/- SC-514, and hypoxic culture +/- SC-514, and PBS + SC-514. Differentially expressed genes (adjusted FDR < 0.05) are indicated in red (up-regulated) or blue (down-regulated). Mean of 3 -5 replicates per experimental condition. **D** Top 10 percentile by FDR-adjusted *P*-value of over-represented genes sets from the GO, KEGG, and REACTOME databases in genes that were significantly up- or down-regulated by treatment with SC-514 alone relative to PBS-injected control HIOs. **E** Venn Diagram showing the number of genes shared between Gene Set I, Gene Set II, and the set of genes that are significantly down-regulated in PBS-injected HIOs treated with SC-514 relative to PBS-injected control HIOs. **F** Top 10 percentile by FDR-adjusted *P*-value of over-represented genes sets from the GO, KEGG, and REACTOME databases in genes that were shared between Gene Set I or II and the set of genes that are significantly down-regulated in HIOs treated with SC-514 relative to PBS-injected control HIOs.

**Figure 5 - Supplement 2.**
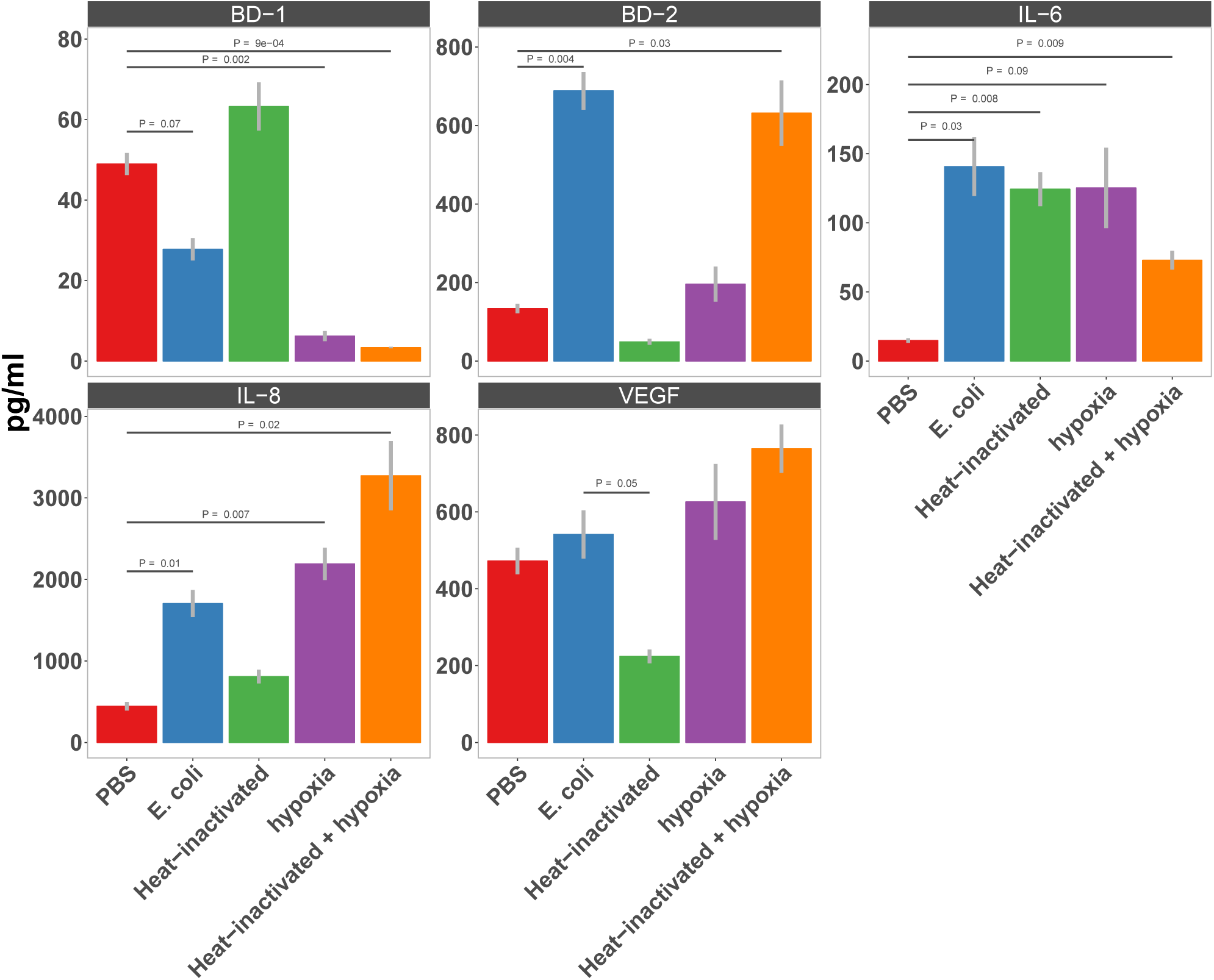
Secretion of AMPs (BD-1 & BD-2), cytokines (IL-6 & IL-8), and the pro-angiogenesis growth factor VEGF in HIOs microinjected with PBS, 10^4^ CFU *E. coli*, or an equivalent concentration of heat-inactivated *E. coli* and cultured under standard cell culture conditions or hypoxic conditions (1% O_2_, 5% CO_2_, 94% N_2_) for 24 h as measured by ELISA. *N* = 7-8 HIOs per condition.

**Figure 6 - Supplement 1.**
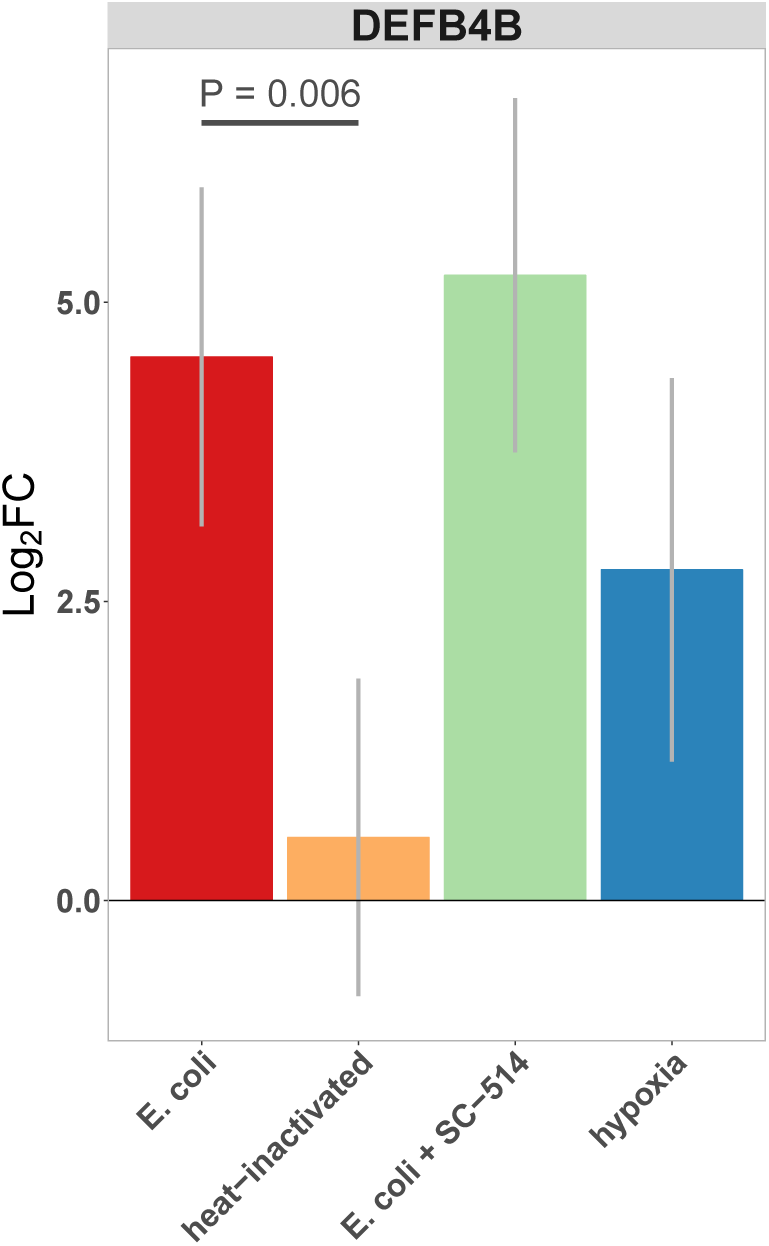
Normalized fold change in expression of DEFB4B, a duplicated gene encoding human *p*-defensin 2 (BD-2) peptide, in each of the conditions indicated relative to PBS control treatment. **C** Concentration of BD-2 peptide in culture supernatant at 24 h as measured by ELISA in HIO cultures treated as indicated. *P* values represents the results of a two-tailed Student’s *t*-test for the comparison indicated.

**Figure 6 - Supplement 2.**
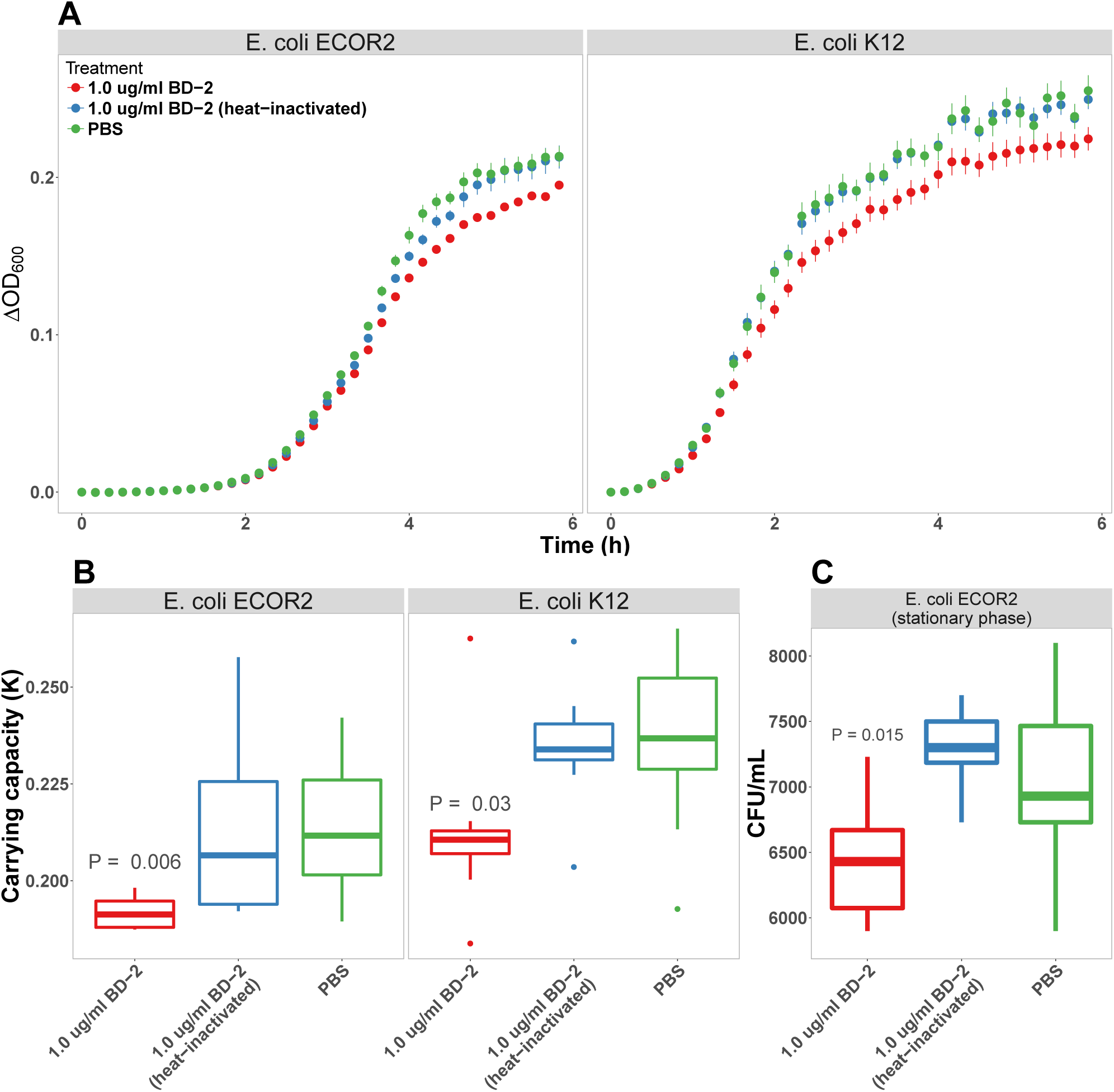
**A** Optical density (600 nm) of *E. coli* str. ECOR2 and *E. coli* str. K12 suspension cultures supplemented with PBS, BD-2 (1 *μ*g/ml), or heat-inactivated BD-2 at 10 min intervals over a 6 h period at 37 °C. **B** Carrying capacity (*к*) of *E. coli* cultures incubated with PBS, BD-2 (1 *μ*g/ml), or heat-inactivated BD-2 during log-phase growth (panel A). *P* values represents the results of a one-tailed Student’s *t*-test relative to PBS treatment. For panels A and B, *N* = 8 per experimental condition for each *E. coli* strain. *C* CFU/ml of late stationary phase *E. coli* str. ECOR2 cultures diluted in PBS and supplemented with BD-2 (1 *μ*g/ml) or heat-inactivated BD-2 for 6 h at 37 °C. *N* = 8 (BD-2 and heat inactivated BD-2) or 16 (PBS).

**Figure 7 - Supplement 1.**
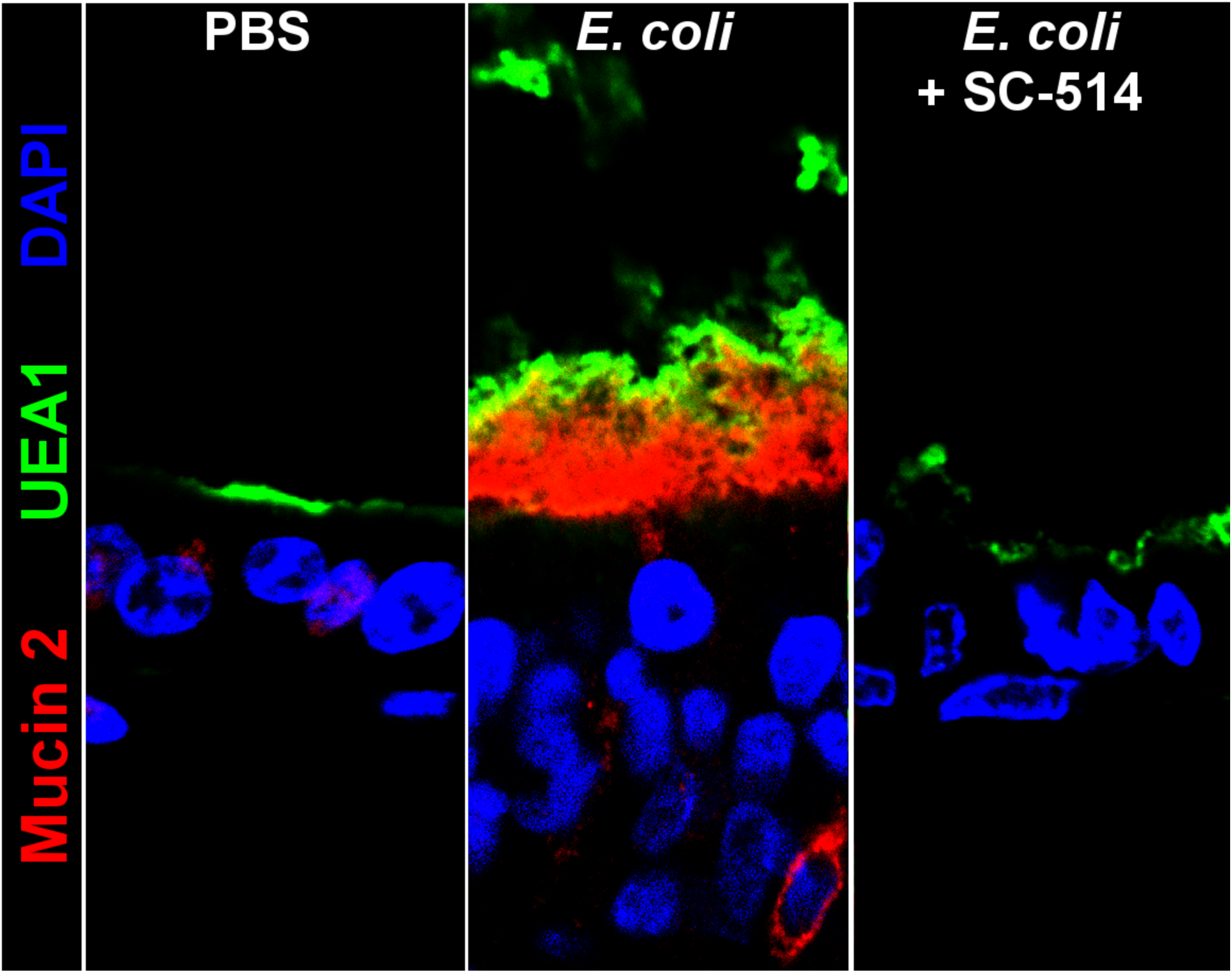
Representative confocal micrographs of HIOs treated as indicated. Fluorescent immunostaining pseudocoloring applied as indicated in the figure legend. 40X optical magnification with 3X digital zoom.

**Figure 8 - Supplement 1.**
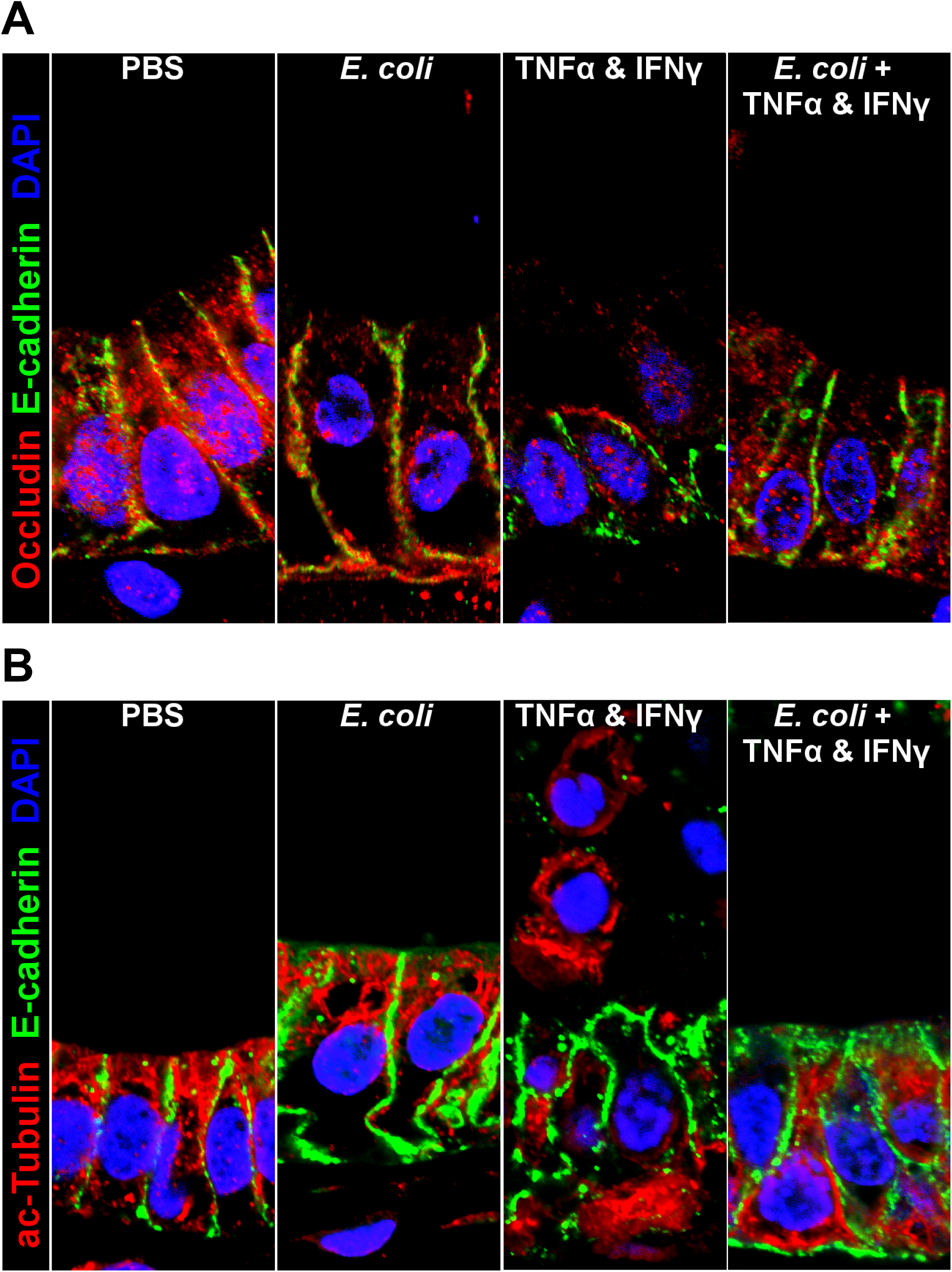
Representative confocal micrographs of HIOs treated as indicated. Fluorescent immunostaining pseudocoloring applied as indicated in the figure legends. 60X optical magnification with 2X digital zoom.

## References

Aagaard K, Ma J, Antony KM, Ganu R, Petrosino J, Versalovic J. The placenta harbors a unique microbiome. Sci Transl Med. 2014 May; 6(237):237ra65. doi: 10.1126/scitranslmed.3008599.

Abrahamsson TR, Jakobsson HE, Andersson AF, Björkstén B, Engstrand L, Jenmalm MC. Low gut microbiota diversity in early infancy precedes asthma at school age. Clin Exp Allergy. 2014 Jun; 44(6):842–50. doi: 10.1111/cea.12253.

Abrams GD, Bauer H, Sprinz H. Influence of the normal flora on mucosal morphology and cellular renewal in the ileum. A comparison of germ-free and conventional mice. Lab Invest. 1963 Mar; 12:355–64.

Afrazi A, Branca MF, Sodhi CP, Good M, Yamaguchi Y, Egan CE, Lu P, Jia H, Shaffiey S, Lin J, Ma C, Vincent G, Prindle T Jr, Weyandt S, Neal MD, Ozolek JA, Wiersch J, Tschurtschenthaler M, Shiota C, Gittes GK, et al. Toll-like receptor 4-mediated endoplasmic reticulum stress in intestinal crypts induces necrotizing enterocolitis. J Biol Chem. 2014 Apr; 289(14):9584–99. doi: 10.1074/jbc.M113.526517.

Ahmad R, Sorrell MF, Batra SK, Dhawan P, Singh AB. Gut permeability and mucosal inflammation: bad, good or context dependent. Mucosal Immunol. 2017 Mar; 10(2):307–317. doi: 10.1038/mi.2016.128.

Ahn DH, Crawley SC, Hokari R, Kato S, Yang SC, Li JD, Kim YS. TNF-alpha activates MUC2 transcription via NF-kappaB but inhibits via JNK activation. Cell Physiol Biochem. 2005; 15(1-4):29–40. doi: 10.1159/000083636.

Albenberg L, Esipova TV, Judge CP, Bittinger K, Chen J, Laughlin A, Grunberg S, Baldassano RN, Lewis JD, Li H, Thom SR, Bushman FD, Vinogradov SA, Wu GD. Correlation between intraluminal oxygen gradient and radial partitioning of intestinal microbiota. Gastroenterology. 2014 Nov; 147(5):1055–63.e8. doi: 10.1053/j.gastro.2014.07.020.

Alfaleh K, Anabrees J, Bassler D, Al-Kharfi T. Probiotics for prevention of necrotizing enterocolitis in preterm infants. Cochrane Database Syst Rev. 2011 Mar; (3):CD005496. doi: 10.1002/14651858.CD005496.pub3.

Arias-Loste MT, Fábrega E, López-Hoyos M, Crespo J. The Crosstalk between Hypoxia and Innate Immunity in the Development of Obesity-Related Nonalcoholic Fatty Liver Disease. Biomed Res Int. 2015; 2015:319745. doi: 10.1155/2015/319745.

Arrieta MC, Stiemsma LT, Dimitriu PA, Thorson L, Russell S, Yurist-Doutsch S, Kuzeljevic B, Gold MJ, Britton HM, Lefebvre DL, Subbarao P, Mandhane P, Becker A, McNagny KM, Sears MR, Kollmann T, CHILD Study Investigators, Mohn WW, Turvey SE, Brett Finlay B. Early infancy microbial and metabolic alterations affect risk of childhood asthma. Sci Transl Med. 2015 Sep; 7(307):307ra152. doi: 10.1126/scitranslmed.aab2271.

Arteel GE, Thurman RG, Raleigh JA. Reductive metabolism of the hypoxia marker pimonidazole is regulated by oxygen tension independent of the pyridine nucleotide redox state. Eur J Biochem. 1998 May; 253(3):743–50.

Artis D. Epithelial-cell recognition of commensal bacteria and maintenance of immune homeostasis in the gut. Nat Rev Immunol. 2008 Jun; 8(6):411–20. doi: 10.1038/nri2316.

Ashburner M, Ball CA, Blake JA, Botstein D, Butler H, Cherry JM, Davis AP, Dolinski K, Dwight SS, Eppig JT, Harris MA, Hill DP, Issel-Tarver L, Kasarskis A, Lewis S, Matese JC, Richardson JE, Ringwald M, Rubin GM, Sherlock G. Gene ontology: tool for the unification of biology. The Gene Ontology Consortium. Nat Genet. 2000 May; 25(1):25–9. doi: 10.1038/75556.

Aurora M, Spence JR. hPSC-derived lung and intestinal organoids as models of human fetal tissue. Dev Biol. 2016 Dec; 420(2):230–238. doi: 10.1016/j.ydbio.2016.06.006.

Bäckhed F, Roswall J, Peng Y, Feng Q, Jia H, Kovatcheva-Datchary P, Li Y, Xia Y, Xie H, Zhong H, Khan MT, Zhang J, Li J, Xiao L, Al-Aama J, Zhang D, Lee YS, Kotowska D, Colding C, Tremaroli V, et al. Dynamics and Stabilization of the Human Gut Microbiome during the First Year of Life. Cell Host Microbe. 2015 Jun; 17(6):852. doi: 10.1016/j.chom.2015.05.012.

Balimane PV, Chong S. Cell culture-based models for intestinal permeability: a critique. Drug Discov Today. 2005 Mar; 10(5):335–43. doi: 10.1016/S1359-6446(04)03354-9.

Bastide P, Darido C, Pannequin J, Kist R, Robine S, Marty-Double C, Bibeau F, Scherer G, Joubert D, Hollande F, Blache P, Jay P. Sox9 regulates cell proliferation and is required for Paneth cell differentiation in the intestinal epithelium. J Cell Biol. 2007 Aug; 178(4):635–48. doi: 10.1083/jcb.200704152.

Bates JM, Mittge E, Kuhlman J, Baden KN, Cheesman SE, Guillemin K. Distinct signals from the microbiota promote different aspects of zebrafish gut differentiation. Dev Biol. 2006 Sep; 297(2):374–86. doi: 10.1016/j.ydbio.2006.05.006.

Bergstrom KS, Xia L. Mucin-type O-glycans and their roles in intestinal homeostasis. Glycobiology. 2013 Sep; 23(9):1026–37. doi: 10.1093/glycob/cwt045.

Bermudez-Brito M, Plaza-Díaz J, Fontana L, Muñoz-Quezada S, Gil A. In vitro cell and tissue models for studying host-microbe interactions: a review. Br J Nutr. 2013 Jan; 109 Suppl 2:S27–34. doi: 10.1017/S0007114512004023.

Bevins CL, Salzman NH. Paneth cells, antimicrobial peptides and maintenance of intestinal homeostasis. Nat Rev Microbiol. 2011 May; 9(5):356–68. doi: 10.1038/nrmicro2546.

Bischoff SC, Barbara G, Buurman W, Ockhuizen T, Schulzke JD, Serino M, Tilg H, Watson A, Wells JM. Intestinal permeability–a new target for disease prevention and therapy. BMC Gastroenterol. 2014 Nov; 14:189. doi: 10.1186/s12876-014-0189-7.

Borre YE, O’Keeffe GW, Clarke G, Stanton C, Dinan TG, Cryan JF. Microbiota and neurodevelopmental windows: implications for brain disorders. Trends Mol Med. 2014 Sep; 20(9):509–18. doi: 10.1016/j.molmed.2014.05.002.

Bray NL, Pimentel H, Melsted P, Pachter L. Near-optimal probabilistic RNA-seq quantification. Nat Biotechnol. 2016 May; 34(5):525–7. doi: 10.1038/nbt.3519.

Broderick NA, Buchon N, Lemaitre B. Microbiota-induced changes in drosophila melanogaster host gene expression and gut morphology. MBio. 2014; 5(3):e01117–14. doi: 10.1128/mBio.01117-14.

Brogden KA. Antimicrobial peptides: pore formers or metabolic inhibitors in bacteria? Nat Rev Microbiol. 2005 Mar; 3(3):238–50. doi: 10.1038/nrmicro1098.

Bry L, Falk PG, Midtvedt T, Gordon JI. A model of host-microbial interactions in an open mammalian ecosystem. Science. 1996 Sep; 273(5280):1380–3.

Buffie CG, Pamer EG. Microbiota-mediated colonization resistance against intestinal pathogens. Nat Rev Immunol. 2013 Nov; 13(11):790–801. doi: 10.1038/nri3535.

Buisine MP, Desreumaux P, Leteurtre E, Copin MC, Colombel JF, Porchet N, Aubert JP. Mucin gene expression in intestinal epithelial cells in Crohn’s disease. Gut. 2001 Oct; 49(4):544–51.

Buisine MP, Devisme L, Savidge TC, Gespach C, Gosselin B, Porchet N, Aubert JP. Mucin gene expression in human embryonic and fetal intestine. Gut. 1998 Oct; 43(4):519–24.

Cash HL, Whitham CV, Behrendt CL, Hooper LV. Symbiotic bacteria direct expression of an intestinal bactericidal lectin. Science. 2006 Aug; 313(5790):1126–30. doi: 10.1126/science.1127119.

Cheesman SE, Neal JT, Mittge E, Seredick BM, Guillemin K. Epithelial cell proliferation in the developing zebrafish intestine is regulated by the Wnt pathway and microbial signaling via Myd88. Proc Natl Acad Sci U S A. 2011 Mar; 108 Suppl 1:4570–7. doi: 10.1073/pnas.1000072107.

Chin AM, Hill DR, Aurora M, Spence JR. Morphogenesis and maturation of the embryonic and postnatal intestine. Semin Cell Dev Biol. 2017 Feb; doi: 10.1016/j.semcdb.2017.01.011.

Cho I, Yamanishi S, Cox L, Methé BA, Zavadil J, Li K, Gao Z, Mahana D, Raju K, Teitler I, Li H, Alekseyenko AV, Blaser MJ. Antibiotics in early life alter the murine colonic microbiome and adiposity. Nature. 2012 Aug; 488(7413):621–6. doi: 10.1038/nature11400.

Clarke G, O’Mahony SM, Dinan TG, Cryan JF. Priming for health: gut microbiota acquired in early life regulates physiology, brain and behaviour. Acta Paediatr. 2014 Aug; 103(8):812–9. doi: 10.1111/apa.12674.

Colgan SP, Curtis VF, Campbell EL. The inflammatory tissue microenvironment in IBD. Inflamm Bowel Dis. 2013 Sep; 19(10):2238–44. doi: 10.1097/MIB.0b013e31828dcaaf.

Collado MC, Rautava S, Aakko J, Isolauri E, Salminen S. Human gut colonisation may be initiated in utero by distinct microbial communities in the placenta and amniotic fluid. Sci Rep. 2016 Mar; 6:23129. doi: 10.1038/srep23129.

Cornick S, Tawiah A, Chadee K. Roles and regulation of the mucus barrier in the gut. Tissue Barriers. 2015; 3(1-2):e982426. doi: 10.4161/21688370.2014.982426.

Costello EK, Stagaman K, Dethlefsen L, Bohannan BJ, Relman DA. The application of ecological theory toward an understanding of the human microbiome. Science. 2012 Jun; 336(6086):1255–62. doi: 10.1126/science.1224203.

Croft D, Mundo AF, Haw R, Milacic M, Weiser J, Wu G, Caudy M, Garapati P, Gillespie M, Kamdar MR, Jassal B, Jupe S, Matthews L, May B, Palatnik S, Rothfels K, Shamovsky V, Song H, Williams M, Birney E, et al. The Reactome pathway knowledgebase. Nucleic Acids Res. 2014 Jan; 42(Database issue):D472–7. doi: 10.1093/nar/gkt1102.

Cullen TW, Schofield WB, Barry NA, Putnam EE, Rundell EA, Trent MS, Degnan PH, Booth CJ, Yu H, Goodman AL. Gut microbiota. Antimicrobial peptide resistance mediates resilience of prominent gut commensals during inflammation. Science. 2015 Jan; 347(6218):170–5. doi: 10.1126/science.1260580.

Dedhia PH, Bertaux-Skeirik N, Zavros Y, Spence JR. Organoid Models of Human Gastrointestinal Development and Disease. Gastroenterology. 2016 05; 150(5):1098–112. doi: 10.1053/j.gastro.2015.12.042.

Desai MS, Seekatz AM, Koropatkin NM, Kamada N, Hickey CA, Wolter M, Pudlo NA, Kitamoto S, Terrapon N, Muller A, Young VB, Henrissat B, Wilmes P, Stappenbeck TS, Núñez G, Martens EC. A Dietary Fiber-Deprived Gut Microbiota Degrades the Colonic Mucus Barrier and Enhances Pathogen Susceptibility. Cell. 2016 Nov; 167(5):1339–1353.e21. doi: 10.1016/j.cell.2016.10.043.

Desbonnet L, Clarke G, Shanahan F, Dinan TG, Cryan JF. Microbiota is essential for social development in the mouse. Mol Psychiatry. 2014 Feb; 19(2):146–8. doi: 10.1038/mp.2013.65.

Diaz Heijtz R, Wang S, Anuar F, Qian Y, Björkholm B, Samuelsson A, Hibberd ML, Forssberg H, Pettersson S. Normal gut microbiota modulates brain development and behavior. Proc Natl Acad Sci U S A. 2011 Feb; 108(7):3047–52. doi: 10.1073/pnas.1010529108.

Donaldson GP, Lee SM, Mazmanian SK. Gut biogeography of the bacterial microbiota. Nat Rev Microbiol. 2016 Jan; 14(1):20–32. doi: 10.1038/nrmicro3552.

Dupont A, Heinbockel L, Brandenburg K, Hornef MW. Antimicrobial peptides and the enteric mucus layer act in concert to protect the intestinal mucosa. Gut Microbes. 2014; 5(6):761–5. doi: 10.4161/19490976.2014.972238.

Dye BR, Dedhia PH, Miller AJ, Nagy MS, White ES, Shea LD, Spence JR. A bioengineered niche promotes in vivo engraftment and maturation of pluripotent stem cell derived human lung organoids. Elife. 2016 Sep; 5. doi: 10.7554/eLife.19732.

Einerhand AW, Renes IB, Makkink MK, van der Sluis M, Büller HA, Dekker J. Role of mucins in inflammatory bowel disease: important lessons from experimental models. Eur J Gastroenterol Hepatol. 2002 Jul; 14(7):757–65.

Emami CN, Mittal R, Wang L, Ford HR, Prasadarao NV. Role of neutrophils and macrophages in the pathogenesis of necrotizing enterocolitis caused by Cronobacter sakazakii. J Surg Res. 2012 Jan; 172(1):18–28. doi: 10.1016/j.jss.2011.04.019.

Engevik MA, Yacyshyn MB, Engevik KA,Wang J, Darien B, Hassett DJ, Yacyshyn BR,Worrell RT. Human Clostridium difficile infection: altered mucus production and composition. Am J Physiol Gastrointest Liver Physiol. 2015 Mar; 308(6):G510–24. doi: 10.1152/ajpgi.00091.2014.

Erkosar B, Storelli G, Mitchell M, Bozonnet L, Bozonnet N, Leulier F. Pathogen Virulence Impedes Mutualist-Mediated Enhancement of Host Juvenile Growth via Inhibition of Protein Digestion. Cell Host Microbe. 2015 Oct; 18(4):445–55. doi: 10.1016/j.chom.2015.09.001.

Espey MG. Role of oxygen gradients in shaping redox relationships between the human intestine and its microbiota. Free Radic Biol Med. 2013 Feb; 55:130–40. doi: 10.1016/j.freeradbiomed.2012.10.554.

Fabregat A, Sidiropoulos K, Garapati P, Gillespie M, Hausmann K, Haw R, Jassal B, Jupe S, Korninger F, McKay S, Matthews L, May B, Milacic M, Rothfels K, Shamovsky V,Webber M,Weiser J, Williams M,Wu G, Stein L, et al. The Reactome pathway Knowledgebase. Nucleic Acids Res. 2016 Jan; 44(D1):D481–7. doi: 10.1093/nar/gkv1351.

Fanaro S, Chierici R, Guerrini P, Vigi V. Intestinal microflora in early infancy: composition and development. Acta Paediatr Suppl. 2003 Sep; 91(441):48–55.

Favier CF, Vaughan EE, De Vos WM, Akkermans AD. Molecular monitoring of succession of bacterial communities in human neonates. Appl Environ Microbiol. 2002 Jan; 68(1):219–26.

Finkbeiner SR, Freeman JJ, Wieck MM, El-Nachef W, Altheim CH, Tsai YH, Huang S, Dyal R, White ES, Grikscheit TC, Teitelbaum DH, Spence JR. Generation of tissue-engineered small intestine using embryonic stem cell-derived human intestinal organoids. Biol Open. 2015 Oct; 4(11):1462–72. doi: 10.1242/bio.013235.

Finkbeiner SR, Hill DR, Altheim CH, Dedhia PH, Taylor MJ, Tsai YH, Chin AM, Mahe MM, Watson CL, Freeman JJ, Nattiv R, Thomson M, Klein OD, Shroyer NF, Helmrath MA, Teitelbaum DH, Dempsey PJ, Spence JR. Transcriptome-wide Analysis Reveals Hallmarks of Human Intestine Development and Maturation In Vitro and In Vivo. Stem Cell Reports. 2015 Jun; doi: 10.1016/j.stemcr.2015.04.010.

Finkbeiner SR, Zeng XL, Utama B, Atmar RL, Shroyer NF, Estes MK. Stem cell-derived human intestinal organoids as an infection model for rotaviruses. MBio. 2012; 3(4):e00159–12. doi: 10.1128/mBio.00159-12.

Fisher EM, Khan M, Salisbury R, Kuppusamy P. Noninvasive monitoring of small intestinal oxygen in a rat model of chronic mesenteric ischemia. Cell Biochem Biophys. 2013 Nov; 67(2):451–9. doi: 10.1007/s12013-013-9611-y.

Forbester JL, Goulding D, Vallier L, Hannan N, Hale C, Pickard D, Mukhopadhyay S, Dougan G. Interaction of Salmonella enterica Serovar Typhimurium with Intestinal Organoids Derived from Human Induced Pluripotent Stem Cells. Infect Immun. 2015 Jul; 83(7):2926–34. doi: 10.1128/IAI.00161-15.

Ford H, Watkins S, Reblock K, Rowe M. The role of inflammatory cytokines and nitric oxide in the pathogenesis of necrotizing enterocolitis. J Pediatr Surg. 1997 Feb; 32(2):275–82.

Ford HR, Sorrells DL, Knisely AS. Inflammatory cytokines, nitric oxide, and necrotizing enterocolitis. Semin Pediatr Surg. 1996 Aug; 5(3):155–9.

Forster R, Chiba K, Schaeffer L, Regalado SG, Lai CS, Gao Q, Kiani S, Farin HF, Clevers H, Cost GJ, Chan A, Rebar EJ, Urnov FD, Gregory PD, Pachter L, Jaenisch R, Hockemeyer D. Human intestinal tissue with adult stem cell properties derived from pluripotent stem cells. Stem Cell Reports. 2014 Jun; 2(6):838–52. doi: 10.1016/j.stemcr.2014.05.001.

Furuyama K, Kawaguchi Y, Akiyama H, Horiguchi M, Kodama S, Kuhara T, Hosokawa S, Elbahrawy A, Soeda T, Koizumi M, Masui T, Kawaguchi M, Takaori K, Doi R, Nishi E, Kakinoki R, Deng JM, Behringer RR, Nakamura T, Uemoto S. Continuous cell supply from a Sox9-expressing progenitor zone in adult liver, exocrine pancreas and intestine. Nat Genet. 2011 Jan; 43(1):34–41. doi: 10.1038/ng.722.

Fusunyan RD, Nanthakumar NN, Baldeon ME, Walker WA. Evidence for an innate immune response in the immature human intestine: toll-like receptors on fetal enterocytes. Pediatr Res. 2001 Apr; 49(4):589–93. doi: 10.1203/00006450-200105000-00006.

Ganz T. Defensins: antimicrobial peptides of innate immunity. Nat Rev Immunol. 2003 Sep; 3(9):710–20. doi: 10.1038/nri1180.

García-Lafuente A, Antolín M, Guarner F, Crespo E, Malagelada JR. Modulation of colonic barrier function by the composition of the commensal flora in the rat. Gut. 2001 Apr; 48(4):503–7.

Gene Ontology Consortium. Gene Ontology Consortium: going forward. Nucleic Acids Res. 2015 Jan; 43(Database issue):D1049–56. doi: 10.1093/nar/gku1179.

Gensollen T, Iyer SS, Kasper DL, Blumberg RS. How colonization by microbiota in early life shapes the immune system. Science. 2016 Apr; 352(6285):539–44. doi: 10.1126/science.aad9378.

Gerdes J, Lemke H, Baisch H, Wacker HH, Schwab U, Stein H. Cell cycle analysis of a cell proliferation-associated human nuclear antigen defined by the monoclonal antibody Ki-67. J Immunol. 1984 Oct; 133(4):1710–5.

Gilmore TD. Introduction to NF-kappaB: players, pathways, perspectives. Oncogene. 2006 Oct; 25(51):6680–4. doi: 10.1038/sj.onc.1209954.

Glover LE, Lee JS, Colgan SP. Oxygen metabolism and barrier regulation in the intestinal mucosa. J Clin Invest. 2016 Oct; 126(10):3680–3688. doi: 10.1172/JCI84429.

Gosalbes MJ, Llop S, Vallès Y, Moya A, Ballester F, Francino MP. Meconium microbiota types dominated by lactic acid or enteric bacteria are differentially associated with maternal eczema and respiratory problems in infants. Clin Exp Allergy. 2013 Feb; 43(2):198–211. doi: 10.1111/cea.12063.

Goto Y, Obata T, Kunisawa J, Sato S, Ivanov II, Lamichhane A, Takeyama N, Kamioka M, Sakamoto M, Matsuki T, Setoyama H, Imaoka A, Uematsu S, Akira S, Domino SE, Kulig P, Becher B, Renauld JC, Sasakawa C, Umesaki Y, et al. Innate lymphoid cells regulate intestinal epithelial cell glycosylation. Science. 2014 Sep; 345(6202):1254009. doi: 10.1126/science.1254009.

Greenwood C, Morrow AL, Lagomarcino AJ, Altaye M, Taft DH, Yu Z, Newburg DS, Ward DV, Schibler KR. Early empiric antibiotic use in preterm infants is associated with lower bacterial diversity and higher relative abundance of Enterobacter. J Pediatr. 2014 Jul; 165(1):23–9. doi: 10.1016/j.jpeds.2014.01.010.

Grenz A, Clambey E, Eltzschig HK. Hypoxia signaling during intestinal ischemia and inflammation. Curr Opin Crit Care. 2012 Apr; 18(2):178–85. doi: 10.1097/MCC.0b013e3283514bd0.

Gruette FK, Horn R, Haenel H. Nutrition and Biochemical Micro-Ecological Processes in the Rectum of Infants. Zeitschrift fuür Kinderheilkunde. 1965; 93:28.

Hackam DJ, Good M, Sodhi CP. Mechanisms of gut barrier failure in the pathogenesis of necrotizing enterocolitis: Toll-like receptors throw the switch. Semin Pediatr Surg. 2013 May; 22(2):76–82. doi: 10.1053/j.sempedsurg.2013.01.003.

Halpern MD, Holubec H, Dominguez JA, Meza YG, Williams CS, Ruth MC, McCuskey RS, Dvorak B. Hepatic inflammatory mediators contribute to intestinal damage in necrotizing enterocolitis. Am J Physiol Gastrointest Liver Physiol. 2003 Apr; 284(4):G695–702. doi: 10.1152/ajpgi.00353.2002.

Hansson GC. Role of mucus layers in gut infection and inflammation. Curr Opin Microbiol. 2012 Feb; 15(1):57–62. doi: 10.1016/j.mib.2011.11.002.

Harder J, Siebert R, Zhang Y, Matthiesen P, Christophers E, Schlegelberger B, Schröder JM. Mapping of the gene encoding human beta-defensin-2 (DEFB2) to chromosome region 8p22-p23.1. Genomics. 1997 Dec; 46(3):472–5. doi: 10.1006/geno.1997.5074.

Henry L, Wickham H, Chang W. ggstance: Horizontal ‘ggplot2’ Components; 2016, https://CRAN.R-project.org/package=ggstance, r package version 0.3.

Hill DR. Hill_HIO_Colonization_2017. Github. 2017; 0a563a7. github.com/hilldr/Hill_HIO_Colonization_2017.

Hill DR. HIO_microinjection. Github. 2017; 49ef21c. github.com/hilldr/HIO_microinjection.

Hill DR, Spence JR. Gastrointestinal Organoids: Understanding the Molecular Basis of the Host-Microbe Interface. Cell Mol Gastroenterol Hepatol. 2017 Mar; 3(2):138–149. doi: 10.1016/j.jcmgh.2016.11.007.

Hirota SA, Fines K, Ng J, Traboulsi D, Lee J, Ihara E, Li Y, Willmore WG, Chung D, Scully MM, Louie T, Medlicott S, Lejeune M, Chadee K, Armstrong G, Colgan SP, Muruve DA, MacDonald JA, Beck PL. Hypoxia-inducible factor signaling provides protection in Clostridium difficile-induced intestinal injury. Gastroenterology. 2010 Jul; 139(1):259–69.e3. doi: 10.1053/j.gastro.2010.03.045.

Hooper LV, Xu J, Falk PG, Midtvedt T, Gordon JI. A molecular sensor that allows a gut commensal to control its nutrient foundation in a competitive ecosystem. Proc Natl Acad Sci U S A. 1999 Aug; 96(17):9833–8.

Hviid A, Svanström H, Frisch M. Antibiotic use and inflammatory bowel diseases in childhood. Gut. 2011 Jan; 60(1):49–54. doi: 10.1136/gut.2010.219683.

Ijssennagger N, Belzer C, Hooiveld GJ, Dekker J, van Mil SW, Müller M, Kleerebezem M, van der Meer R. Gut microbiota facilitates dietary heme-induced epithelial hyperproliferation by opening the mucus barrier in colon. Proc Natl Acad Sci U S A. 2015 Aug; 112(32):10038–43. doi: 10.1073/pnas.1507645112.

Jakobsson HE, Rodríguez-Piñeiro AM, Schütte A, Ermund A, Boysen P, Bemark M, Sommer F, Bäckhed F, Hansson GC, Johansson ME. The composition of the gut microbiota shapes the colon mucus barrier. EMBO Rep. 2015 Feb; 16(2):164–77. doi: 10.15252/embr.201439263.

Johansson ME, Hansson GC. Immunological aspects of intestinal mucus and mucins. Nat Rev Immunol. 2016 10; 16(10):639–49. doi: 10.1038/nri.2016.88.

Joly S, Maze C, McCray PB Jr, Guthmiller JM. Human beta-defensins 2 and 3 demonstrate strain-selective activity against oral microorganisms. J Clin Microbiol. 2004 Mar; 42(3):1024–9.

Josephson R, Sykes G, Liu Y, Ording C, Xu W, Zeng X, Shin S, Loring J, Maitra A, Rao MS, Auerbach JM. A molecular scheme for improved characterization of human embryonic stem cell lines. BMC Biol. 2006 Aug; 4:28. doi: 10.1186/1741-7007-4--28.

Kaiko GE, Ryu SH, Koues OI, Collins PL, Solnica-Krezel L, Pearce EJ, Pearce EL, Oltz EM, Stappenbeck TS. The Colonic Crypt Protects Stem Cells from Microbiota-Derived Metabolites. Cell. 2016 Jun; 165(7):1708–20. doi: 10.1016/j.cell.2016.05.018.

Kanehisa M, Goto S. KEGG: kyoto encyclopedia of genes and genomes. Nucleic Acids Res. 2000 Jan; 28(1):27–30.

Kawai T, Akira S. Signaling to NF-kappaB by Toll-like receptors. Trends Mol Med. 2007 Nov; 13(11):460–9. doi: 10.1016/j.molmed.2007.09.002.

Kelly CJ, Zheng L, Campbell EL, Saeedi B, Scholz CC, Bayless AJ, Wilson KE, Glover LE, Kominsky DJ, Magnuson A, Weir TL, Ehrentraut SF, Pickel C, Kuhn KA, Lanis JM, Nguyen V, Taylor CT, Colgan SP. Crosstalk between Microbiota-Derived Short-Chain Fatty Acids and Intestinal Epithelial HIF Augments Tissue Barrier Function. Cell Host Microbe. 2015 May; 17(5):662–71. doi: 10.1016/j.chom.2015.03.005.

Khailova L, Dvorak K, Arganbright KM, Halpern MD, Kinouchi T, Yajima M, Dvorak B. Bifidobacterium bifidum improves intestinal integrity in a rat model of necrotizing enterocolitis. Am J Physiol Gastrointest Liver Physiol. 2009 Nov; 297(5):G940–9.

Kim MS, Bae JW. Spatial disturbances in altered mucosal and luminal gut viromes of diet-induced obese mice. Environ Microbiol. 2016 May; 18(5):1498–510. doi: 10.1111/1462-2920.13182.

Kim YG, Sakamoto K, Seo SU, Pickard JM, Gillilland MG 3rd, Pudlo NA, Hoostal M, Li X, Wang TD, Feehley T, Stefka AT, Schmidt TM, Martens EC, Fukuda S, Inohara N, Nagler CR, Núñez G. Neonatal acquisition of Clostridia species protects against colonization by bacterial pathogens. Science. 2017 Apr; 356(6335):315–319. doi: 10.1126/science.aag2029.

Kim YS, Ho SB. Intestinal goblet cells and mucins in health and disease: recent insights and progress. Curr Gastroenterol Rep. 2010 Oct; 12(5):319–30. doi: 10.1007/s11894-010-0131-2.

Kishore N, Sommers C, Mathialagan S, Guzova J, Yao M, Hauser S, Huynh K, Bonar S, Mielke C, Albee L, Weier R, Graneto M, Hanau C, Perry T, Tripp CS. A selective IKK-2 inhibitor blocks NF-kappa B-dependent gene expression in interleukin-1 beta-stimulated synovial fibroblasts. J Biol Chem. 2003 Aug; 278(35):32861–71. doi: 10.1074/jbc.M211439200.

Kisich KO, Heifets L, Higgins M, Diamond G. Antimycobacterial agent based on mRNA encoding human beta-defensin 2 enables primary macrophages to restrict growth of Mycobacterium tuberculosis. Infect Immun. 2001 Apr; 69(4):2692–9. doi: 10.1128/IAI.69.4.2692-2699.2001.

Koenig JE, Spor A, Scalfone N, Fricker AD, Stombaugh J, Knight R, Angenent LT, Ley RE. Succession of microbial consortia in the developing infant gut microbiome. Proc Natl Acad Sci U S A. 2011 Mar; 108 Suppl 1:4578–85. doi: 10.1073/pnas.1000081107.

Koong AC, Chen EY, Giaccia AJ. Hypoxia causes the activation of nuclear factor kappa B through the phosphorylation of I kappa B alpha on tyrosine residues. Cancer Res. 1994 Mar; 54(6):1425–30.

Kremer N, Philipp EE, Carpentier MC, Brennan CA, Kraemer L, Altura MA, Augustin R, Häsler R, Heath-Heckman EA, Peyer SM, Schwartzman J, Rader BA, Ruby EG, Rosenstiel P, McFall-Ngai MJ. Initial symbiont contact orchestrates host-organ-wide transcriptional changes that prime tissue colonization. Cell Host Microbe. 2013 Aug; 14(2):183–94. doi: 10.1016/j.chom.2013.07.006.

Lebenthal A, Lebenthal E. The ontogeny of the small intestinal epithelium. JPEN J Parenter Enteral Nutr. 1999; 23(5 Suppl):S3–6. doi: 10.1177/014860719902300502.

Leslie JL, Huang S, Opp JS, Nagy MS, Kobayashi M, Young VB, Spence JR. Persistence and toxin production by Clostridium difficile within human intestinal organoids result in disruption of epithelial paracellular barrier function. Infect Immun. 2015 Jan; 83(1):138–45. doi: 10.1128/IAI.02561-14.

Li H, Limenitakis JP, Fuhrer T, Geuking MB, Lawson MA, Wyss M, Brugiroux S, Keller I, Macpherson JA, Rupp S, Stolp B, Stein JV, Stecher B, Sauer U, McCoy KD, Macpherson AJ. The outer mucus layer hosts a distinct intestinal microbial niche. Nat Commun. 2015 Sep; 6:8292. doi: 10.1038/ncomms9292.

Litvak V, Ramsey SA, Rust AG, Zak DE, Kennedy KA, Lampano AE, Nykter M, Shmulevich I, Aderem A. Function of C/EBPdelta in a regulatory circuit that discriminates between transient and persistent TLR4-induced signals. Nat Immunol. 2009 Apr; 10(4):437–43. doi: 10.1038/ni.1721.

Love MI, Huber W, Anders S. Moderated estimation of fold change and dispersion for RNA-seq data with DESeq2. Genome Biology. 2014; 15:550. doi: 10.1186/s13059-014-0550-8.

Luo, Weijun, Brouwer, Cory. Pathview: an R/Bioconductor package for pathway-based data integration and visualization. Bioinformatics. 2013; 29(14):1830–1831. doi: 10.1093/bioinformatics/btt285.

Malago JJ. Contribution of microbiota to the intestinal physicochemical barrier. Benef Microbes. 2015; 6(3):295–311. doi: 10.3920/BM2014.0041.

Marcobal A, Southwick AM, Earle KA, Sonnenburg JL. A refined palate: bacterial consumption of host glycans in the gut. Glycobiology. 2013 Sep; 23(9):1038–46. doi: 10.1093/glycob/cwt040.

Matsuo Y, Nishizaki C, Drexler HG. Efficient DNA fingerprinting method for the identification of cross-culture contamination of cell lines. Hum Cell. 1999 Sep; 12(3):149–54.

McCracken KW, Catá EM, Crawford CM, Sinagoga KL, Schumacher M, Rockich BE, Tsai YH, Mayhew CN, Spence JR, Zavros Y, Wells JM. Modelling human development and disease in pluripotent stem-cell-derived gastric organoids. Nature. 2014 Dec; 516(7531):400–4. doi: 10.1038/nature13863.

McCracken KW, Howell JC, Wells JM, Spence JR. Generating human intestinal tissue from pluripotent stem cells in vitro. Nat Protoc. 2011 Nov; 6(12):1920–8. doi: 10.1038/nprot.2011.410.

McFall-Ngai MJ. The importance of microbes in animal development: lessons from the squid-vibrio symbiosis. Annu Rev Microbiol. 2014; 68:177–94. doi: 10.1146/annurev-micro-091313-103654.

McLoughlin K, Schluter J, Rakoff-Nahoum S, Smith AL, Foster KR. Host Selection of Microbiota via Differential Adhesion. Cell Host Microbe. 2016 Apr; 19(4):550–9. doi: 10.1016/j.chom.2016.02.021.

Michielan A, D’Incà R. Intestinal Permeability in Inflammatory Bowel Disease: Pathogenesis, Clinical Evaluation, and Therapy of Leaky Gut. Mediators Inflamm. 2015; 2015:628157. doi: 10.1155/2015/628157.

Miyoshi H, Ajima R, Luo CT, Yamaguchi TP, Stappenbeck TS. Wnt5a potentiates TGF-beta signaling to promote colonic crypt regeneration after tissue injury. Science. 2012 Oct; 338(6103):108–13. doi: 10.1126/science. 1223821.

Miyoshi H, Stappenbeck TS. In vitro expansion and genetic modification of gastrointestinal stem cells in spheroid culture. Nat Protoc. 2013 Dec; 8 Aug; 133(2):539–46. doi: 10.1053/j.gastro.2007.05.020.

Morrow AL, Lagomarcino AJ, Schibler KR, Taft DH, Yu Z, Wang B, Altaye M, Wagner M, Gevers D, Ward DV, Kennedy MA, Huttenhower C, Newburg DS. Early microbial and metabolomic signatures predict later onset of necrotizing enterocolitis in preterm infants. Microbiome. 2013; 1(1):13. doi: 10.1186/2049-2618-1-13.

Muniz LR, Knosp C, Yeretssian G. Intestinal antimicrobial peptides during homeostasis, infection, and disease. Front Immunol. 2012; 3:310. doi: 10.3389/fimmu.2012.00310.

Ménard S, Förster V, Lotz M, Gütle D, Duerr CU, Gallo RL, Henriques-Normark B, Pütsep K, Andersson M, Glocker EO, Hornef MW. Developmental switch of intestinal antimicrobial peptide expression. J Exp Med. 2008 Jan; 205(1):183–93. doi: 10.1084/jem.20071022.

Nanthakumar N, Meng D, Goldstein AM, Zhu W, Lu L, Uauy R, Llanos A, Claud EC, Walker WA. The mechanism of excessive intestinal inflammation in necrotizing enterocolitis: an immature innate immune response. PLoS One. 2011; 6(3):e17776. doi: 10.1371/journal.pone.0017776.

Neal JT, Peterson TS, Kent ML, Guillemin K. H. pylori virulence factor CagA increases intestinal cell proliferation by Wnt pathway activation in a transgenic zebrafish model. Dis Model Mech. 2013 May; 6(3):802–10. doi: 10.1242/dmm.011163.

Neu J. Gastrointestinal maturation and implications for infant feeding. Early Hum Dev. 2007 Dec; 83(12):767–75. doi: 10.1016/j.earlhumdev.2007.09.009.

Neu J, Walker WA. Necrotizing enterocolitis. N Engl J Med. 2011 Jan; 364(3):255–64. doi: 10.1056/NEJMra1005408.

Nguyen TL, Vieira-Silva S, Liston A, Raes J. How informative is the mouse for human gut microbiota research? Dis Model Mech. 2015 Jan; 8(1):1–16. doi: 10.1242/dmm.017400.

Nishimura E, Eto A, Kato M, Hashizume S, Imai S, Nisizawa T, Hanada N. Oral streptococci exhibit diverse susceptibility to human beta-defensin-2: antimicrobial effects of hBD-2 on oral streptococci. Curr Microbiol. 2004 Feb; 48(2):85–7. doi: 10.1007/s00284-003-4108-3.

Ochman H, Selander RK. Standard reference strains of Escherichia coli from natural populations. J Bacteriol. 1984 Feb; 157(2):690–3.

Oliver KM, Taylor CT, Cummins EP. Hypoxia. Regulation of NFkappaB signalling during inflammation: the role of hydroxylases. Arthritis Res Ther. 2009; 11(1):215. doi: 10.1186/ar2575.

O’Neil DA. Regulation of expression of beta-defensins: endogenous enteric peptide antibiotics. Mol Immunol. 2003 Nov; 40(7):445–50.

Ostaff MJ, Stange EF, Wehkamp J. Antimicrobial peptides and gut microbiota in homeostasis and pathology. EMBO Mol Med. 2013 10; 5(10):1465–83. doi: 10.1002/emmm.201201773.

Palmer C, Bik EM, DiGiulio DB, Relman DA, Brown PO. Development of the human infant intestinal microbiota. PLoS Biol. 2007 Jul; 5(7):e177. doi: 10.1371/journal.pbio.0050177.

Peterson LW, Artis D. Intestinal epithelial cells: regulators of barrier function and immune homeostasis. Nat Rev Immunol. 2014 Mar; 14(3):141–53. doi: 10.1038/nri3608.

R Core Team. R: A Language and Environment for Statistical Computing. R Foundation for Statistical Computing, Vienna, Austria; 2017, https://www.R-project.org/.

Renz H, Brandtzaeg P, Hornef M. The impact of perinatal immune development on mucosal homeostasis and chronic inflammation. Nat Rev Immunol. 2011 Dec; 12(1):9–23. doi: 10.1038/nri3112.

Renz H, Brandtzaeg P, Hornef M. The impact of perinatal immune development on mucosal homeostasis and chronic inflammation. Nat Rev Immunol. 2012 Jan; 12(1):9–23. doi: 10.1038/nri3112.

Rhee SJ, Walker WA, Cherayil BJ. Developmentally regulated intestinal expression of IFN-gamma and its target genes and the age-specific response to enteric Salmonella infection. J Immunol. 2005 Jul; 175(2):1127–36.

Rius J, Guma M, Schachtrup C, Akassoglou K, Zinkernagel AS, Nizet V, Johnson RS, Haddad GG, Karin M. NFkappaB links innate immunity to the hypoxic response through transcriptional regulation of HIF-1alpha. Nature. 2008 Jun; 453(7196):807–11. doi: 10.1038/nature06905.

Rivera-Chávez F, Zhang LF, Faber F, Lopez CA, Byndloss MX, Olsan EE, Xu G, Velazquez EM, Lebrilla CB, Winter SE, Bäumler AJ. Depletion of Butyrate-Producing Clostridia from the Gut Microbiota Drives an Aerobic Luminal Expansion of Salmonella. Cell Host Microbe. 2016 Apr; 19(4):443–54. doi: 10.1016/j.chom.2016.03.004.

Robinson J. Cochrane in context: probiotics for prevention of necrotizing enterocolitis in preterm infants. Evid Based Child Health. 2014 Sep; 9(3):672–4. doi: 10.1002/ebch.1977.

Rodriguez-Pineiro AM, Bergstrom JH, Ermund A, Gustafsson JK, Schutte A, Johansson ME, Hansson GC. Studies of mucus in mouse stomach, small intestine, and colon. II. Gastrointestinal mucus proteome reveals Muc2 and Muc5ac accompanied by a set of core proteins. Am J Physiol Gastrointest Liver Physiol. 2013 Sep; 305(5):G348–56. doi: 10.1152/ajpgi.00047.2013.

Rolig AS, Parthasarathy R, Burns AR, Bohannan BJ, Guillemin K. Individual Members of the Microbiota Disproportionately Modulate Host Innate Immune Responses. Cell Host Microbe. 2015 Nov; 18(5):613–20. doi: 10.1016/j.chom.2015.10.009.

Round JL, Mazmanian SK. The gut microbiota shapes intestinal immune responses during health and disease. Nat Rev Immunol. 2009 May; 9(5):313–23. doi: 10.1038/nri2515.

Salzman NH, Ghosh D, Huttner KM, Paterson Y, Bevins CL. Protection against enteric salmonellosis in transgenic mice expressing a human intestinal defensin. Nature. 2003 Apr; 422(6931):522–6. doi: 10.1038/nature01520.

Salzman NH, Hung K, Haribhai D, Chu H, Karlsson-Sjöberg J, Amir E, Teggatz P, Barman M, Hayward M, Eastwood D, Stoel M, Zhou Y, Sodergren E, Weinstock GM, Bevins CL, Williams CB, Bos NA. Enteric defensins are essential regulators of intestinal microbial ecology. Nat Immunol. 2010 Jan; 11(1):76–83. doi: 10.1038/ni.1825.

Sander LE, Davis MJ, Boekschoten MV, Amsen D, Dascher CC, Ryffel B, Swanson JA, Müller M, Blander JM. Detection of prokaryotic mRNA signifies microbial viability and promotes immunity. Nature. 2011 May; 474(7351):385–9. doi: 10.1038/nature10072.

Sanderson IR, Walker WA. Development of the Gastrointestinal Tract. No. v. 1 in Development of the Gastrointestinal Tract, B.C. Decker; 2000. https://books.google.com/books?id=YhgKZ_dvda0C.

Schluter J, Foster KR. The evolution of mutualism in gut microbiota via host epithelial selection. PLoS Biol. 2012; 10(11):e1001424. doi: 10.1371/journal.pbio.1001424.

Schmidt TM, Kao JY. A little O2 may go a long way in structuring the GI microbiome. Gastroenterology. 2014 Nov; 147(5):956–9. doi: 10.1053/j.gastro.2014.09.025.

Shaw SY, Blanchard JF, Bernstein CN. Association between the use of antibiotics in the first year of life and pediatric inflammatory bowel disease. Am J Gastroenterol. 2010 Dec; 105(12):2687–92. doi: 10.1038/ajg.2010.398.

Shawki A, McCole DF. Mechanisms of Intestinal Epithelial Barrier Dysfunction by Adherent-Invasive Escherichia coli. Cell Mol Gastroenterol Hepatol. 2017 Jan; 3(1):41–50. doi: 10.1016/j.jcmgh.2016.10.004.

Shreiner AB, Kao JY, Young VB. The gut microbiome in health and in disease. Curr Opin Gastroenterol. 2015 Jan; 31(1):69–75. doi: 10.1097/MOG.0000000000000139.

Sommer F, Nookaew I, Sommer N, Fogelstrand P, Bäckhed F. Site-specific programming of the host epithelial transcriptome by the gut microbiota. Genome Biol. 2015; 16:62. doi: 10.1186/s13059-015-0614-4.

Spence JR, Mayhew CN, Rankin SA, Kuhar MF, Vallance JE, Tolle K, Hoskins EE, Kalinichenko VV, Wells SI, Zorn AM, Shroyer NF, Wells JM. Directed differentiation of human pluripotent stem cells into intestinal tissue in vitro. Nature. 2011 Feb; 470(7332):105–9. doi: 10.1038/nature09691.

Stallman RM. EMACS the Extensible, Customizable Self-documenting Display Editor. ACM SIGOA Newsletter. 1981 Apr; 2(1-2):147–156. http://doi.acm.org/10.1145/1159890.806466, doi: 10.1145/1159890.806466.

Subramanian A, Tamayo P, Mootha VK, Mukherjee S, Ebert BL, Gillette MA, Paulovich A, Pomeroy SL, Golub TR, Lander ES, Mesirov JP. Gene set enrichment analysis: a knowledge-based approach for interpreting genomewide expression profiles. Proc Natl Acad Sci U S A. 2005 Oct; 102(43):15545–50. doi: 10.1073/pnas.0506580102.

Suenaert P, Bulteel V, Lemmens L, Noman M, Geypens B, Van Assche G, Geboes K, Ceuppens JL, Rutgeerts P. Anti-tumor necrosis factor treatment restores the gut barrier in Crohn’s disease. Am J Gastroenterol. 2002 Aug; 97(8):2000–4. doi: 10.1111/j.1572-0241.2002.05914.x.

Tan X, Hsueh W, Gonzalez-Crussi F. Cellular localization of tumor necrosis factor (TNF)-alpha transcripts in normal bowel and in necrotizing enterocolitis. TNF gene expression by Paneth cells, intestinal eosinophils, and macrophages. Am J Pathol. 1993 Jun; 142(6):1858–65.

Tanner SM, Berryhill TF, Ellenburg JL, Jilling T, Cleveland DS, Lorenz RG, Martin CA. Pathogenesis of necrotizing enterocolitis: modeling the innate immune response. Am J Pathol. 2015 Jan; 185(1):4–16. doi: 10.1016/j.ajpath.2014.08.028.

Tsai YH, Nattiv R, Dedhia PH, Nagy MS, Chin AM, Thomson M, Klein OD, Spence JR. In vitro patterning of pluripotent stem cell-derived intestine recapitulates in vivo human development. Development. 2017 Mar; 144(6):1045–1055. doi: 10.1242/dev.138453.

Tsutsumi-Ishii Y, Nagaoka I. NF-kappa B-mediated transcriptional regulation of human beta-defensin-2 gene following lipopolysaccharide stimulation. J Leukoc Biol. 2002 Jan; 71(1):154–62.

Turner JR. Intestinal mucosal barrier function in health and disease. Nat Rev Immunol. 2009 Nov; 9(11):799–809. doi: 10.1038/nri2653.

Underwood MA, Arriola J, Gerber CW, Kaveti A, Kalanetra KM, Kananurak A, Bevins CL, Mills DA, Dvorak B. Bifidobacterium longum subsp. infantis in experimental necrotizing enterocolitis: alterations in inflammation, innate immune response, and the microbiota. Pediatr Res. 2014 Oct; 76(4):326–33. doi: 10.1038/pr.2014.102.

Upperman JS, Potoka D, Grishin A, Hackam D, Zamora R, Ford HR. Mechanisms of nitric oxide-mediated intestinal barrier failure in necrotizing enterocolitis. Semin Pediatr Surg. 2005 Aug; 14(3):159–66. doi: 10.1053/j.sempedsurg.2005.05.004.

Vaishnava S, Behrendt CL, Ismail AS, Eckmann L, Hooper LV. Paneth cells directly sense gut commensals and maintain homeostasis at the intestinal host-microbial interface. Proc Natl Acad Sci U S A. 2008 Dec; 105(52):20858–63. doi: 10.1073/pnas.0808723105.

Varki A. Essentials of glycobiology. Cold Spring Harbor, New York: Cold Spring Harbor Laboratory Press; 2017.

Veereman-Wauters G. Neonatal gut development and postnatal adaptation. Eur J Pediatr. 1996 Aug; 155(8):627–32.

Vora P, Youdim A, Thomas LS, Fukata M, Tesfay SY, Lukasek K, Michelsen KS, Wada A, Hirayama T, Arditi M, Abreu MT. Beta-defensin-2 expression is regulated by TLR signaling in intestinal epithelial cells. J Immunol. 2004 Nov; 173(9):5398–405.

Wang F, Graham WV, Wang Y, Witkowski ED, Schwarz BT, Turner JR. Interferon-gamma and tumor necrosis factor-alpha synergize to induce intestinal epithelial barrier dysfunction by up-regulating myosin light chain kinase expression. Am J Pathol. 2005 Feb; 166(2):409–19.

Wang F, Schwarz BT, Graham WV, Wang Y, Su L, Clayburgh DR, Abraham C, Turner JR. IFN-gamma-induced TNFR2 expression is required for TNF-dependent intestinal epithelial barrier dysfunction. Gastroenterology. 2006 Oct; 131(4):1153–63. doi: 10.1053/j.gastro.2006.08.022.

Wang G, Li X, Wang Z. APD3: the antimicrobial peptide database as a tool for research and education. Nucleic Acids Res. 2016 Jan; 44(D1):D1087–93. doi: 10.1093/nar/gkv1278.

Ward DV, Scholz M, Zolfo M, Taft DH, Schibler KR, Tett A, Segata N, Morrow AL. Metagenomic Sequencing with Strain-Level Resolution Implicates Uropathogenic E. coli in Necrotizing Enterocolitis and Mortality in Preterm Infants. Cell Rep. 2016 Mar; 14(12):2912–24. doi: 10.1016/j.celrep.2016.03.015.

Watson CL, Mahe MM, Múnera J, Howell JC, Sundaram N, Poling HM, Schweitzer JI, Vallance JE, Mayhew CN, Sun Y, Grabowski G, Finkbeiner SR, Spence JR, Shroyer NF, Wells JM, Helmrath MA. An in vivo model of human small intestine using pluripotent stem cells. Nat Med. 2014 Nov; 20(11):1310–4. doi: 10.1038/nm.3737.

Wattam AR, Davis JJ, Assaf R, Boisvert S, Brettin T, Bun C, Conrad N, Dietrich EM, Disz T, Gabbard JL, Gerdes S, Henry CS, Kenyon RW, Machi D, Mao C, Nordberg EK, Olsen GJ, Murphy-Olson DE, Olson R, Overbeek R, et al. Improvements to PATRIC, the all-bacterial Bioinformatics Database and Analysis Resource Center. Nucleic Acids Res. 2017 Jan; 45(D1):D535–D542. doi: 10.1093/nar/gkw1017.

Wickham H. ggplot2: Elegant Graphics for Data Analysis. Springer-Verlag New York; 2009. http://ggplot2.org.

Wopereis H, Oozeer R, Knipping K, Belzer C, Knol J. The first thousand days - intestinal microbiology of early life: establishing a symbiosis. Pediatr Allergy Immunol. 2014 Aug; 25(5):428–38. doi: 10.1111/pai.12232.

Wullaert A, Bonnet MC, Pasparakis M. NF-kappaB in the regulation of epithelial homeostasis and inflammation. Cell Res. 2011 Jan; 21(1):146–58. doi: 10.1038/cr.2010.175.

Xiao C, Ghosh S. NF-kappaB, an evolutionarily conserved mediator of immune and inflammatory responses. Adv Exp Med Biol. 2005; 560:41–5. doi: 10.1007/0-387-24180-9_5.

Xie L, Xue X, Taylor M, Ramakrishnan SK, Nagaoka K, Hao C, Gonzalez FJ, Shah YM. Hypoxia-inducible factor/MAZ-dependent induction of caveolin-1 regulates colon permeability through suppression of occludin, leading to hypoxia-induced inflammation. Mol Cell Biol. 2014 Aug; 34(16):3013–23. doi: 10.1128/MCB.00324-14.

Yassour M, Vatanen T, Siljander H, Hämäläinen AM, Härkönen T, Ryhänen SJ, Franzosa EA, Vlamakis H, Huttenhower C, Gevers D, Lander ES, Knip M, DIABIMMUNE Study Group, Xavier RJ. Natural history of the infant gut microbiome and impact of antibiotic treatment on bacterial strain diversity and stability. Sci Transl Med. 2016 Jun; 8(343):343ra81. doi: 10.1126/scitranslmed.aad0917.

Yu G, He QY. ReactomePA: an R/Bioconductor package for reactome pathway analysis and visualization. Molecular BioSystems. 2016; 12(12):477–479. http://pubs.rsc.org/en/Content/ArticleLanding/2015/MB/C5MB00663E, doi: 10.1039/C5MB00663E.

Yu G, Wang LG, Han Y, He QY. clusterProfiler: an R package for comparing biological themes among gene clusters. OMICS: A Journal of Integrative Biology. 2012; 16(5):284–287. doi: 10.1089/omi.2011.0118.

Zeitouni NE, Chotikatum S, von Köckritz-Blickwede M, Naim HY. The impact of hypoxia on intestinal epithelial cell functions: consequences for invasion by bacterial pathogens. Mol Cell Pediatr. 2016 Dec; 3(1):14. doi: 10.1186/s40348-016-0041-y.

Zhang G, Ghosh S. Toll-like receptor-mediated NF-kappaB activation: a phylogenetically conserved paradigm in innate immunity. J Clin Invest. 2001 Jan; 107(1):13–9. doi: 10.1172/JCI11837.

Zheng L, Kelly CJ, Colgan SP. Physiologic hypoxia and oxygen homeostasis in the healthy intestine. A Review in the Theme: Cellular Responses to Hypoxia. Am J Physiol Cell Physiol. 2015 Sep; 309(6):C350–60. doi: 10.1152/ajpcell.00191.2015.

## References

Ochman H, Selander RK. Standard reference strains of Escherichia coli from natural populations. J Bacteriol. 1984 Feb; 157(2):690-3.

Wattam AR, Davis JJ, Assaf R, Boisvert S, Brettin T, Bun C, Conrad N, Dietrich EM, Disz T, Gabbard JL, Gerdes S, Henry CS, Kenyon RW, Machi D, Mao C, Nordberg EK, Olsen GJ, Murphy-Olson DE, Olson R, Overbeek R, et al. Improvements to PATRIC, the all-bacterial Bioinformatics Database and Analysis Resource Center. Nucleic Acids Res. 2017 Jan; 45(D1):D535-D542. doi: 10.1093/nar/gkw1017.

